# Stress granules and protein aggregates reveal intracellular resource competition

**DOI:** 10.1101/2025.11.08.687377

**Authors:** Hannah E. Buchholz, Sean A. Martin, Jane E. Dorweiler, Derek C. Prosser, Emily M. Sontag, Anita L. Manogaran

## Abstract

Stress granules are biomolecular condensates that form in response to environmental stress and disassemble once normal conditions are restored. However, when disassembly fails, stress granules can persist and solidify. While stress granule solidification has been well documented, the cellular mechanisms underlying the transition from reversible to persistent stress granules remain unclear. Persistent stress granules can seed the formation of pathological aggregates, such as TDP-43 in amyotrophic lateral sclerosis^1, 2^. Although amyloid and tau aggregates are hallmarks of Alzheimer’s disease, a subset of patients also develop TDP-43 deposits, suggesting a possible role for stress granule solidification in Alzheimer’s disease progression^3–5^. Despite theoretical models explaining why persistence and ensuing solidification occurs, strong *in vivo* evidence is lacking^6^. Here we show that competition for limited chaperone resources drive stress granule persistence. In the presence of TDP-43 aggregates or yeast amyloid proteins called prions, stress granule disassembly is slowed or halted disassembly. Using yeast prions as a model, we show that the addition of chaperones, specifically the AAA+ ATPase molecular chaperone, Hsp104, resulted in resumption of stress granule disassembly. Our results demonstrate that the competition for shared resources, such as molecular chaperones, can limit stress granule disassembly. We suspect that the presence of pathological aggregates results in resource competition within the aging brain, contributing to the persistence of stress granules and their subsequent solidification and aggregation.

## Main

Aberrant clearance of stress granules is linked with disease such as amyotrophic lateral sclerosis (ALS)^7–9^. Although the mechanisms underlying stress granule persistence are not well understood, several processes have been proposed to contribute to this phenomenon, including protein oxidation that promotes localized liquid-to-solid transitions within the stress granule and the generation of pathological TDP-43 aggregates^1, 2, 8, 10^. However, clinical observations indicate that additional factors are likely involved. TDP-43 aggregates have been identified in fifty-seven percent of post-mortem Alzheimer’s patient brains, along with the characteristic amyloid-ß or tau amyloids^3, 4, 11^. In fact, the presence of TDP-43 deposits is correlated with an amplification of Alzheimer’s symptoms in patients^12^. While TDP-43 aggregation in these patients may be due to stress granule solidification^2^, the trigger of solidification is poorly understood.

Stress granules are composed of RNA-binding proteins and RNAs that assemble into condensates in response to a variety of stresses from glucose starvation to heat treatment. When stress subsides, stress granules quickly disassemble and release protein and RNA back into the cellular environment^13^. While disassembly can be mediated by ubiquitination and post-translational modifications^14–16^, molecular chaperones play a central role in the disassembly process. *in vitro* studies show that the yeast J-domain protein, Sis1, and the Hsp70 protein, Ssa1, work together with the Hsp104 disaggregase to mediate disassembly through a partial threading mechanism that disperses individual stress granule components^17, 18^. These same molecular chaperones are also essential for disassembly *in vivo* since loss or depletion of these chaperones leads to stress granule persistence^19–21^. However, other mechanisms have been also proposed that explain stress granule persistence such as sequestration of essential proteins, impairment of other proteostasis machinery, and competition for resources that normally assist protein folding and disaggregation^22–24^.

Resource competition is well documented in ecology where two different species compete for a resource in limited supply^25^. Resource competition can also occur in cells. The presence of disease-associated protein aggregates depletes the available heat shock cognate protein 70 chaperone resulting in dysfunction of clathrin-mediated endocytosis^26^, indicating that aggregates can sequester and limit chaperone availability for common cellular functions. Since molecular chaperones play an important role in limiting protein aggregation, the presence of one aggregate could divert chaperone resources, resulting in the formation of another aggregate formed through liquid-to-solid phase transitions. Here, intracellular co-existence of two different protein aggregates reveals disproportionate competition for the same molecular chaperones. Hsp104 and co-chaperones, Sis1 and Ssa1, play a central role in disassembling stress granules^19–21^, and fragmenting some yeast prions into smaller transmissible seeds for propagation through mitosis^27–31^. Our work shows that while stress granules generally disassemble quickly *in vivo*, stress granule disassembly is slowed or even halted in cells that contain TDP-43 aggregates or yeast prions. Introduction of excess Hsp104 restores stress granule disassembly kinetics in the presence of prions. Our studies provide a model that suggests the efficient use of chaperone resources by one aggregate could limit the availability of these resources for stress granule clearance, thus resulting in stress granule persistence over time. This persistence could mediate the liquid to solid-like transition within the stress granule to form pathogenic aggregates.

### TDP-43 aggregates limit stress granule disassembly

We are interested in how stress granule disassembly is impacted by the presence of human TDP-43 aggregates. In yeast, common laboratory strains such as BY4741 are used to study stress granule dynamics. However, we found that the 74D-694 strain, commonly used to study yeast aggregating proteins called prions, is more sensitive to acute heat shock (Figure S1A). 74D-694 readily forms stress granules in response to a 42°C heat shock for 45 minutes, visualized using the Pab1-GFP stress granule marker. Upon the release from heat shock, we closely monitored disassembly over five hours by surveying the percentage of cells that contained visible Pab1-GFP stress granules. Stress granules in the 74D-694 background disassembled more gradually over five hours after heat shock release relative to BY4741 (Figure S1B), which provides the advantage of being able to monitor and assess temporal or incremental changes in stress granule disassembly over time.

Using the 74D-694 strain, we wanted to assess how the presence of TDP-43 impacted stress granule disassembly. Induction of a galactose inducible YFP tagged TDP-43 (TDP-43-YFP) results in cellular toxicity and the formation of large irreversible protein aggregates in yeast^32^. However, growth in non-inducing 2% raffinose media results in no toxicity, extremely low level TDP-43 expression, and diffuse cytoplasmic YFP fluorescence (Figure S1C-D). While both TDP-43-YFP and Pab1-mCherry puncta were detected after heat shock in 2% raffinose media (Figure 1A and B, S1D), these puncta were not always colocalized (Figure 1A and S1D). These data indicate that low level expression of TDP-43 results in heat responsive aggregation, similar to stress induced aggregation of cytoplasmic TDP-43 in humans^8, 33^, yet these aggregates do not appear to consistently assemble with Pab1-mCherry stress granules. Next, persistence of TDP-43-YFP visible puncta were monitored over time after heat shock under raffinose conditions. Five hours after heat shock showed only 28.4% of cells contained TDP-43-YFP puncta indicating that these TDP-43 stress responsive puncta were reversible (Figure 1C). However, Pab1-mCherry puncta recovered slower in cells expressing TDP-43-YFP under raffinose conditions compared to empty vector controls (Figure 1D), suggesting that despite TDP-43-YFP forming stress responsive puncta that are dynamic, the presence of TDP-43-YFP has a negative impact on Pab1-mCherry stress granule disassembly kinetics.

**Figure 1.**
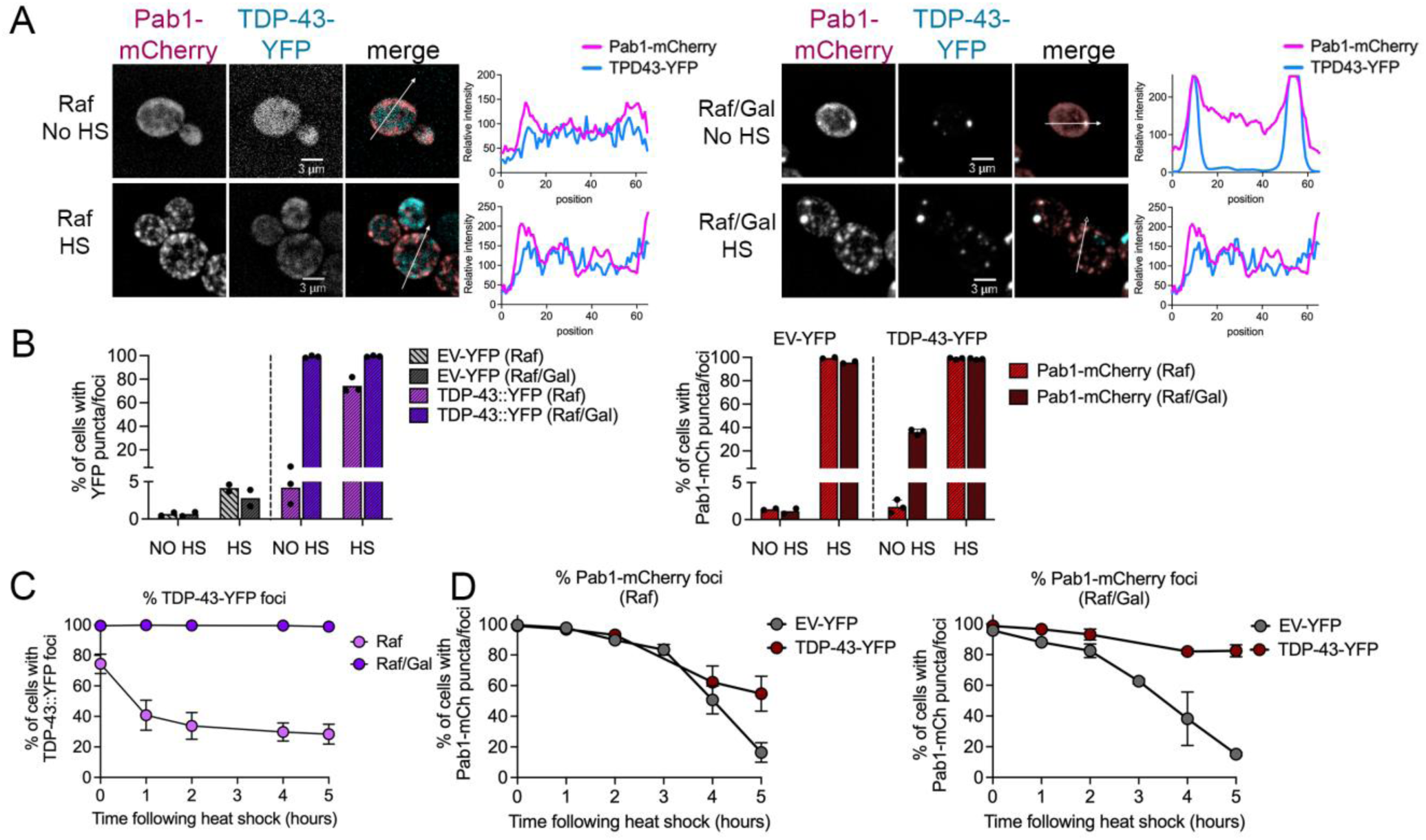
Pab1-mCherry stress granules do not recover in the presence of TDP-43 aggregates. A. Spinning disk confocal microscopy images of strains expressing both TDP-43-YFP and Pab1-mCherry in 2% raffinose (Raf, left) or 2%raffinose/0.1% galactose (Raf/Gal, right) media either untreated (No HS) or immediately after heat shock at 42°C for 45 minutes. Line plots graph the relative fluorescent intensity along the line drawn in the merged image. B. *Left*. Strains containing empty vector (EV-YFP) or TDP-43-YFP were grown in Raf or Raf/Gal media and assessed for the percentage of cells with YFP foci under no heat shock or immediately after heat shock conditions. *Right*. Strains in B were also transformed with Pab1-mCherry and were assessed for the presence of Pab1-mCherry puncta and/or foci. Approximately 200-300 cells were analyzed per trial. C. After release from heat shock, cells in B were assessed for the percentage of cells with TDP-43-YFP foci over time in the indicated media. D. Cells in B were grown in raffinose media (left) of raffinose/galactose media (right) and assessed for the percentage of cells that contain Pab1-mCherry puncta and/or foci after heat shock. Note that the Pab1-mCherry after heat shock contained both large foci and small puncta in A (Raf/Gal). Approximately 200-300 cells were quantified per replicate (n=3) and timepoint.

Induction of TDP-43-YFP expression with 2% galactose results in large visible foci and is associated with toxicity^32^. While growth on 0.1% galactose was not associated with toxicity, TDP-43 expression was significantly increased (Figure S1C), large cytoplasmic foci were detected in the absence of heat shock, and Pab1-mCherry appeared to colocalize with these prominent TDP-43-YFP foci (Figure 1A and S1D). Following heat shock, large Pab1-mCherry still co-localized with large TDP-43-YFP foci, and additional small Pab1-mCherry puncta were detected, which are consistent with the formation of stress granules (Figure 1A and S1C). Despite the loss of TDP-43-YFP aggregates five hours after heat shock in raffinose cultures (Figure 1C), both TDP-43-YFP and Pab1-mCherry puncta and foci persisted after 5 hours in 0.1% galactose (Figure 1C and D). These data suggest that the presence of TDP-43-YFP aggregates limit Pab1-mCherry stress granule recovery.

### Stress granule disassembly is delayed in the presence of yeast prions

While TDP-43-YFP studies revealed impacts on stress granule disassembly, TDP-43 is heterologous in nature and can complicate the understanding of stress granule dynamics in yeast. Therefore, we turned to endogenous yeast aggregates called prions. Two yeast proteins that can form prions are Sup35 and Rnq1, which form [*PSI*^+^] and [*PIN^+^*], respectively^34–36^. We first sought to determine if these two proteins were stress responsive like TDP-43 and Pab1. An integrated GFP-tagged Sup35 where GFP is placed between the N-terminal and middle domain (sup35::N-GFP-MC) or a plasmid containing a C-terminally tagged Rnq1 (Rnq1-GFP) showed that the stress-responsive Sup35 forms small visible puncta in response to transient heat shock, but the Rnq1 protein did not (Figure S2).

Next, we assessed whether stress granule disassembly kinetics were impacted in the absence and presence of a prion. After release of heat shock, Pab1-GFP exhibited normal disassembly over five hours in strains that do not contain a prion (Figure 2A). Yeast prions can exist as different strains or variants, where different aggregate conformations of the same protein result in distinct heritable phenotypes and variable amounts of soluble protein not contained within the aggregate^37, 38^. In weak or strong [*PSI*^+^] variants^34^, stress granule disassembly was delayed. In the case of strong [*PSI*^+^], almost no disassembly was observed, with 94.1% of cells still containing Pab1-GFP puncta after five hours. Strains containing either weak [*PSI*^+^] or two different variants of [*PIN^+^*]^39^ showed delays in stress granule disassembly, with ∼35-50% of cells retaining puncta after 5 hours (Figure 2A). However, stress granules in these populations were able to resolve faster than strong [*PSI^+^*].

**Figure 2.**
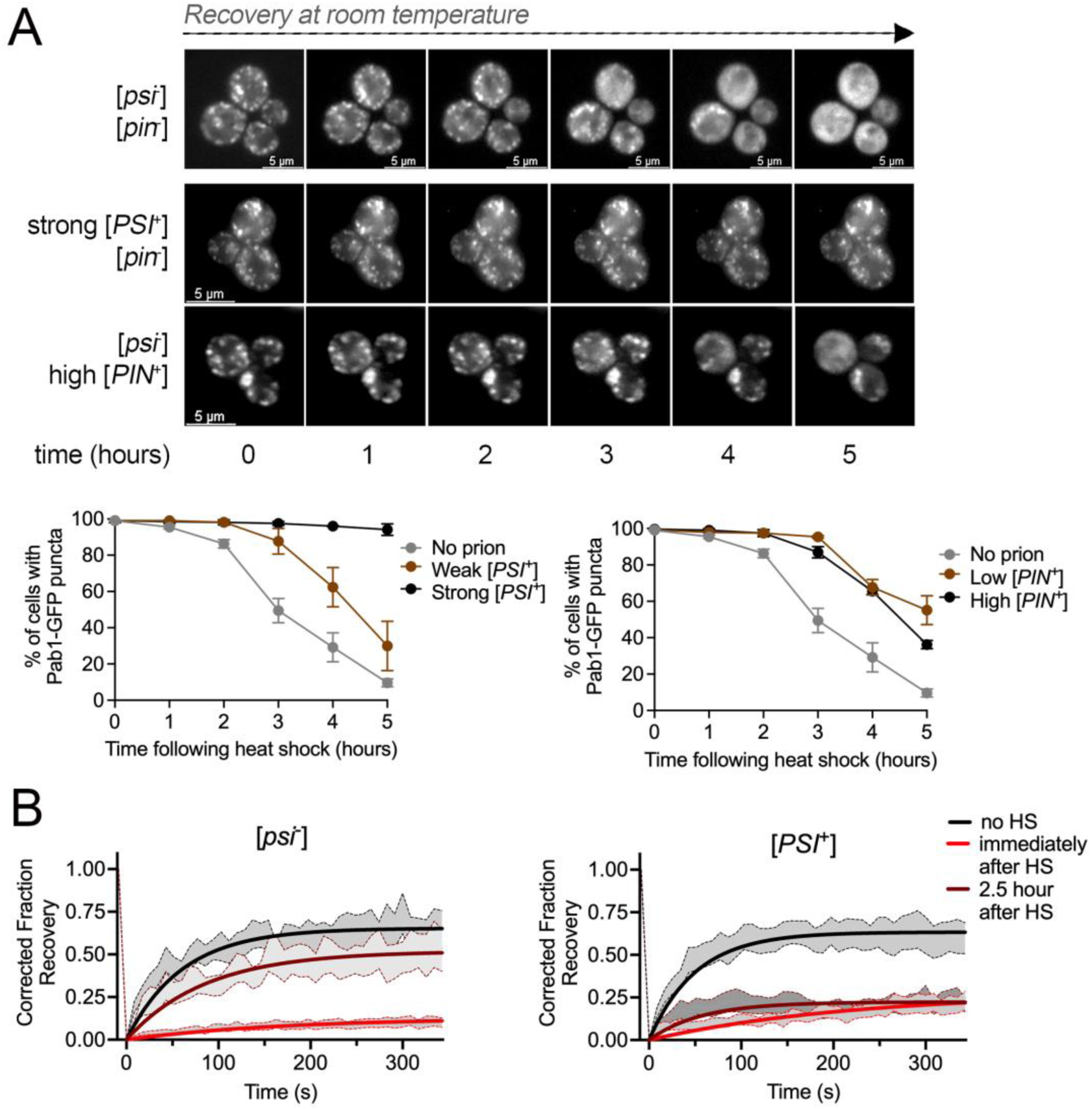
Stress granule disassembly is delayed in the presence of prions. A. *Top,* [*psi*^-^][*pin*^-^], strong [*PSI*^+^], and high [*PIN*^+^] strains containing Pab1-GFP were heat shocked and then imaged by timelapse 3D widefield microscopy at room temperature over five hours. *Bottom*. The indicated strains were heat shocked and were incubated at room temperature. Images were taken of populations at the indicated time points, and the percentage of cells containing visible Pab1-GFP puncta was scored. Approximately 200-300 cells were counted per replicate and timepoint (n=3, mean ±SD). B. FRAP assays of Pab1-GFP puncta in [*psi*^-^] or strong [*PSI*^+^] strains in untreated (no HS, black), immediately after heat shock (red), or 2.5 hours after heat shock (dark red) were imaged over 342 seconds. Data are shown as ±SEM and non-linear regression lines.

To evaluate the dynamic properties of Pab1-GFP, we used fluorescence recovery after photobleaching (FRAP), which provides information about the rate of protein mobility. Under no heat shock conditions, FRAP analysis showed fast Pab1-GFP recovery profiles in strains with and without prions, suggesting that under no heat shock conditions Pab1-GFP is mobile regardless of the prion state (Figure 2B, S3, Table S1 and 2). However, immediately after heat shock, very little recovery was observed, indicating that heat shock results in a loss of Pab1-GFP mobility. Interestingly, differences were observed 2.5 hours after heat shock. [*psi*^-^] cells showed substantial fluorescence recovery, yet [*PSI*^+^] showed almost no fluorescence recovery with approximately 77.8% of Pab1-GFP in the immobile fraction (Figure 2B and Table S1). Taken together, these results indicate that the presence of a prion negatively impacts stress granule disassembly, and in the case of strong [*PSI*^+^], suggest that Pab1 and possibly other stress granule components remain immobile several hours after heat shock.

### Pab1 and Sup35 are colocalized after heat shock

To begin to understand the relationship between Pab1 and prion proteins immediately after heat shock, we performed co-localization studies. Pab1-mCherry was introduced into the endogenously tagged sup35::N-GFP-MC strain. Under no prion conditions, both proteins are diffuse in untreated cells (Figure S4A). While both proteins form puncta in heat shocked cells (Figure 3A and S4A), not all puncta colocalize, as shown by differences in Pearson’s correlation coefficient (PCC) compared to Pab1-GFP/Pab1-mCherry colocalization controls (Figure 3A). In the presence of [*PSI*^+^], non-heat shocked cells showed Pab1-mCherry as diffuse. In contrast, sup35::N-GFP-MC displayed very small puncta (Figure S4B), which were previously reported as fast-moving puncta^40^. Upon heat shock, the distinct sup35::N-GFP-MC puncta are larger and static (Figure S4B) suggesting that heat shock changes the motility and size of sup35::N-GFP-MC aggregates in [*PSI*^+^] cells. These puncta displayed line plot intensity profiles similar to those of Pab1-mCherry, and the corresponding PCC values between channels were similar to those of the Pab1-mCherry/Pab1-GFP positive control, consistent with colocalization of the two proteins (Figure 3A). To further confirm colocalization, the increased resolution of near-TIRF microscopy shows that the majority of Pab1-mCherry puncta have similar patterns to sup35::N-GFP-MC puncta, however there are occasions where no overlap is observed (Figure 3A). Overall, these data suggest that heat shock not only changes the Sup35 prion aggregate, but it also implies that Pab1-mCherry stress granules possibly join these inclusions. In contrast, when assessing aggregation in [*PIN*^+^] strains containing both Rnq1-GFP and Pab1-mCherry, Rnq1-GFP is found in puncta that look similar before and after heat shock, whereas Pab1-mCherry puncta formed after heat shock but do not colocalize with Rnq1-GFP (Figure 3B and S4B). Taken together, our data suggest that these two different yeast prions can associate with heat-induced stress granule components differently.

**Figure 3.**
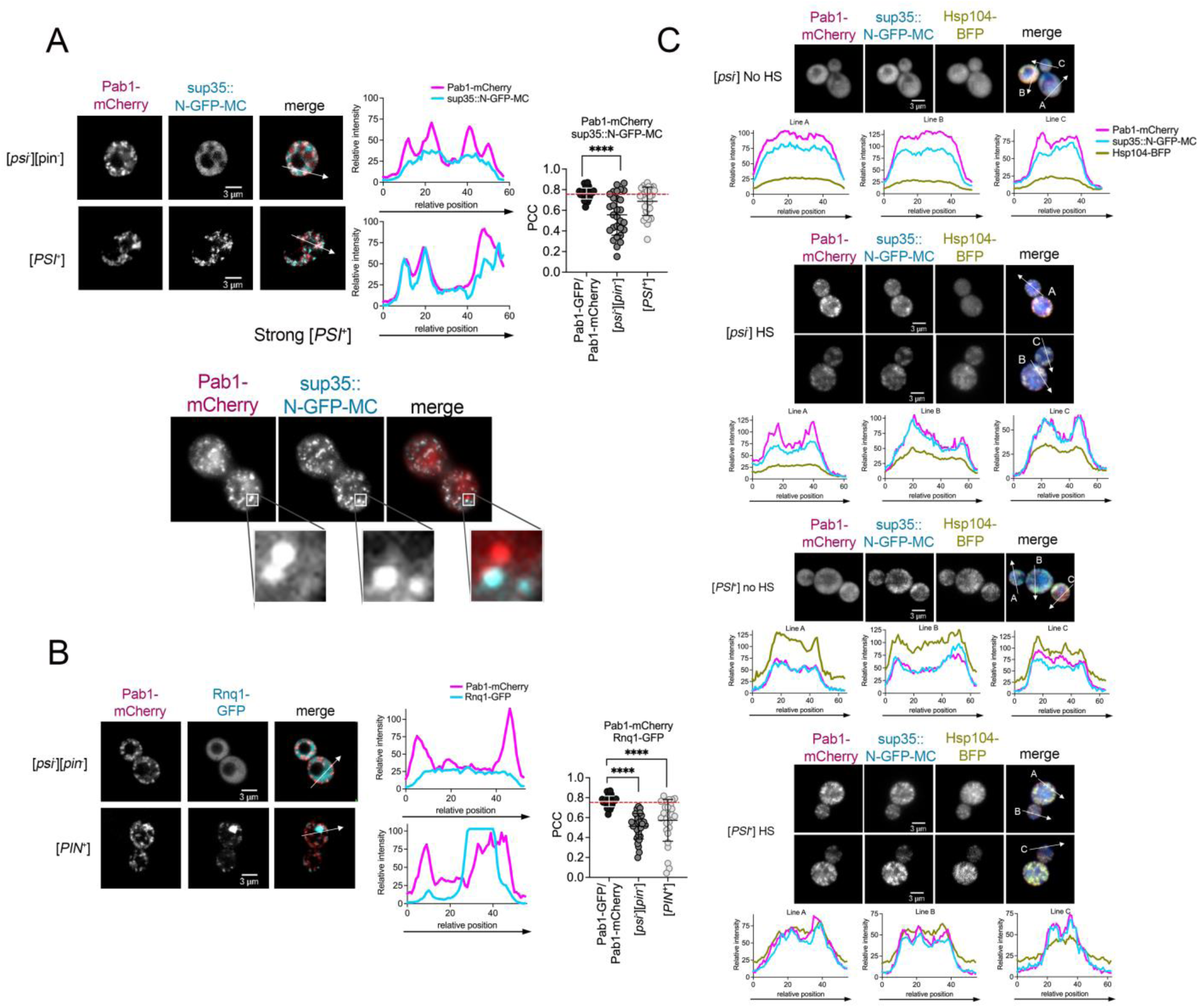
Pab1-mCherry colocalizes with sup35::N-GFP-MC and Hsp104-BFP immediately after heat shock in [*PSI*^+^] cells. A. Strains without ([*psi*^-^][*pin*^-^]) or with the [*PSI*^+^] (strong [*PSI*^+^]) prion were imaged by confocal spinning disk microscopy immediately after heat shock. *Left.* Images of Pab1-mCherry, sup35::N-GFP-MC, and merged images are shown. *Middle.* Line plots are intensity profiles derived from the region of interest indicated by the arrow. *Right*. PCC analysis of colocalization determined for each indicated protein. The dotted line represents the mean positive control PCC between Pab1-GFP and Pab1-mCh (0.763). Experimental PCC means of Pab1-mCherry or sup35::N-GFP-MC were compared to the control mean by two-way ANOVA with Dunnett’s multiple comparisons test (****p≤0.0001) n=30. *Bottom*, Near TIRF imaging of the same strain. Inset shows close-up of areas with and without co-localized signals. B. Confocal imaging, line plots, and PCC values of Pab1-mCherry and Rnq1-GFP in [*PIN*^+^] (high [*PIN*^+^]) strains similar to A. C. [*psi*^-^] or [*PSI*^+^] sup35::N-GFP-MC strains expressing an integrated ADH-Hsp104-BFP and plasmid Pab1-mCherry. Strains were either left untreated (No HS) or imaged immediately after heat shock. Line plots are derived from the arrows. All images are shown at 630X magnification as single Z-planes.

Since heat shock has a direct effect on the heat shock response (HSR), we next asked whether the prions by themselves initiate a HSR or elevated chaperone levels. A HSE-YFP reporter gene, which contains tandem heat shock elements fused to YFP^41^ was introduced into no prion ([*psi*^-^][*pin*^-^]), strong [*PSI*^+^] and high [*PIN*^+^] strains to indirectly measure the heat shock response. While no prion and high [*PIN*^+^] strains have similar HSR outputs, strong [*PSI*^+^] has elevated HSR under no heat shock conditions, which continue after heat shock (Figure S5A). However, all chaperone levels are similar between no prion and prion strains, except for Hsp104, which shows a slight increase in [*PIN*^+^] strains compared to [*PSI*^+^] strains (Figure S5B).

Since Hsp104 has been shown to interact with prion aggregates^42, 43^ and stress granules^44^, we asked whether Hsp104 is colocalized with Pab1-GFP and Sup35 before and after heat shock. Using a [*psi^-^*] strain with a constitutively expressed integrated Hsp104-BFP, we found that Pab1-mCherry, sup35::N-GFP-MC, and Hsp104-BFP are diffuse under no heat shock conditions, and all form aggregates upon heat shock (Figure 3C). Due to the low brightness of the blue fluorescent protein associated with Hsp104-BFP, the aggregates under heat shock were sometimes difficult to clearly observe compared to Pab1-mCherry or sup35::N-GFP-MC. Endogenously tagged Hsp104-GFP confirmed that Hsp104 formed puncta overlapping with Pab1-mCherry stress granules upon heat shock (Figure S6). In the presence of [*PSI*^+^] and no heat shock, small sup35::N-GFP-MC and Hsp104-BFP puncta appear to be colocalized (Figure 3C), consistent with previous studies^42, 45^. Upon heat shock of [*PSI*^+^] cells, sup35::N-GFP-MC puncta appear larger and most colocalize with Pab1-mCherry, similar to our observations in Figure 3A. Hsp104-BFP also form puncta with similar aggregation patterns as the other two proteins (Figure 3C). These similar patterns were confirmed using Pab1-mCherry and Hsp104-GFP (Figure S6). Together, these results suggest that Hsp104 is initially associated with [*PSI*^+^] aggregates before heat shock, and Hsp104 continues to associate with aggregates immediately after heat shock.

### Prions are maintained over time after heat shock despite aberrant stress granule disassembly

Since both prion fragmentation and stress granule disassembly depend upon the same molecular chaperones, it is possible that both aggregates are competing for the same chaperone resources. The ecological theory of resource competition states that a species that uses limiting resources more efficiently will survive over one that requires more of the same resource^25^. If both prions and stress granules compete for chaperones, then based on resource competition theory we would predict that differential efficiency in resource use will allow one aggregate to dominate access to the limiting resource over the other aggregate. For example, if yeast prions utilize chaperones more efficiently than stress granules, then we would expect that the prions continue to be fragmented for propagation while the stress granules have impaired disassembly. To test our hypothesis, we assessed the prion state immediately after heat shock and five hours later. Well trap and SDD-AGE assays of both [*PSI*^+^] and [*PIN*^+^] strains show that the prions are present at both time points (Figure S7). In addition, a phenotypic color assay also confirms that strong [*PSI*^+^] is maintained in the population several days after heat shock (Figure S7A). Based on these results, prion fragmentation is maintained after heat shock despite aberrant stress granule disassembly.

### Hsp104 overexpression relieves Pab1-GFP disassembly delays in prion strains

Since chaperones play a central role in both fragmentation and disassembly, it is possible that the addition of excess chaperone may provide sufficient resources to restore stress granule disassembly in the presence of a prion. To ensure that our system is working, we verified stress granule disassembly kinetics in no prion strains where chaperones were either deleted or overexpressed. As expected, stress granule delays were observed in Hsp104 or Ssa1 deletion strains or Sis1 depletion strains (Figure S8A)^19–21^. Using a plasmid that increases Hsp104 steady state levels by four-fold ^46^, we found that excess Hsp104 slightly delays stress granule disassembly kinetics (Figure S8B). However, introduction of a constitutively expressed Ssa1, which increases Ssa1 levels only by 25% (Figure S8B), completely prevents stress granule disassembly. Interestingly, co-overexpression of Ssa1 and Hsp104 restores disassembly kinetics to near wildtype levels (Figure S8B). It is possible that restoration of proper chaperone stoichiometric balance or heat shock response could result in efficient stress granule disassembly. Introduction of constitutively expressed Sis1, which has eight-fold more protein than wildtype cells, appeared to have no impact on stress granule disassembly (Figure S8B).

We then tested stress granule disassembly kinetics in the presence of [*PSI^+^*] or [*PIN*^+^] with the overexpression of specific chaperones. If stress granules were competing with prions for chaperone resources, then the introduction of excess chaperones should restore stress granule recovery. In the presence of [*PSI*^+^], introduction of excess Sis1 or Ssa1 still resulted in substantial stress granule disassembly delays (Figure S8C and D). In contrast, overexpression of Hsp104 alone showed improved stress granule disassembly resulting in just 35.0% of [*PSI*^+^] cells retaining the Pab1-GFP puncta after 5 hours (Figure 4A), compared to [*PSI*^+^] cells where no chaperone was overexpressed. Despite the disassembly of stress granules by 5 hours with Hsp104 overexpression, phenotypic and biochemical data show that the [*PSI*^+^] prion in the presence of Hsp104 overexpression is maintained in the population (Figure S7A). Together, these data indicate that supplying excess Hsp104 partially improves stress granule recovery in [*PSI*^+^] cells within 5 hours but does not impact prion propagation. [*PIN*^+^] strains show a slightly different response to chaperone overexpression. Overexpression of Ssa1 appears to inhibit stress granule disassembly, comparable to no prion strains, and overexpression of Sis1 seems to have minimal impacts on disassembly in [*PIN*^+^] strains (Figure S8C and D). In contrast, overexpression of Hsp104 completely restored stress granule disassembly in [*PIN*^+^] cells without impacting the propagation of the prion (Figure 4A). Since [*PIN*^+^] requires less Hsp104 for propagation than [*PSI*^+^] ^47^, it is possible that there is more Hsp104 chaperone available in [*PIN*^+^] strains to mediate stress granule disassembly.

**Figure 4.**
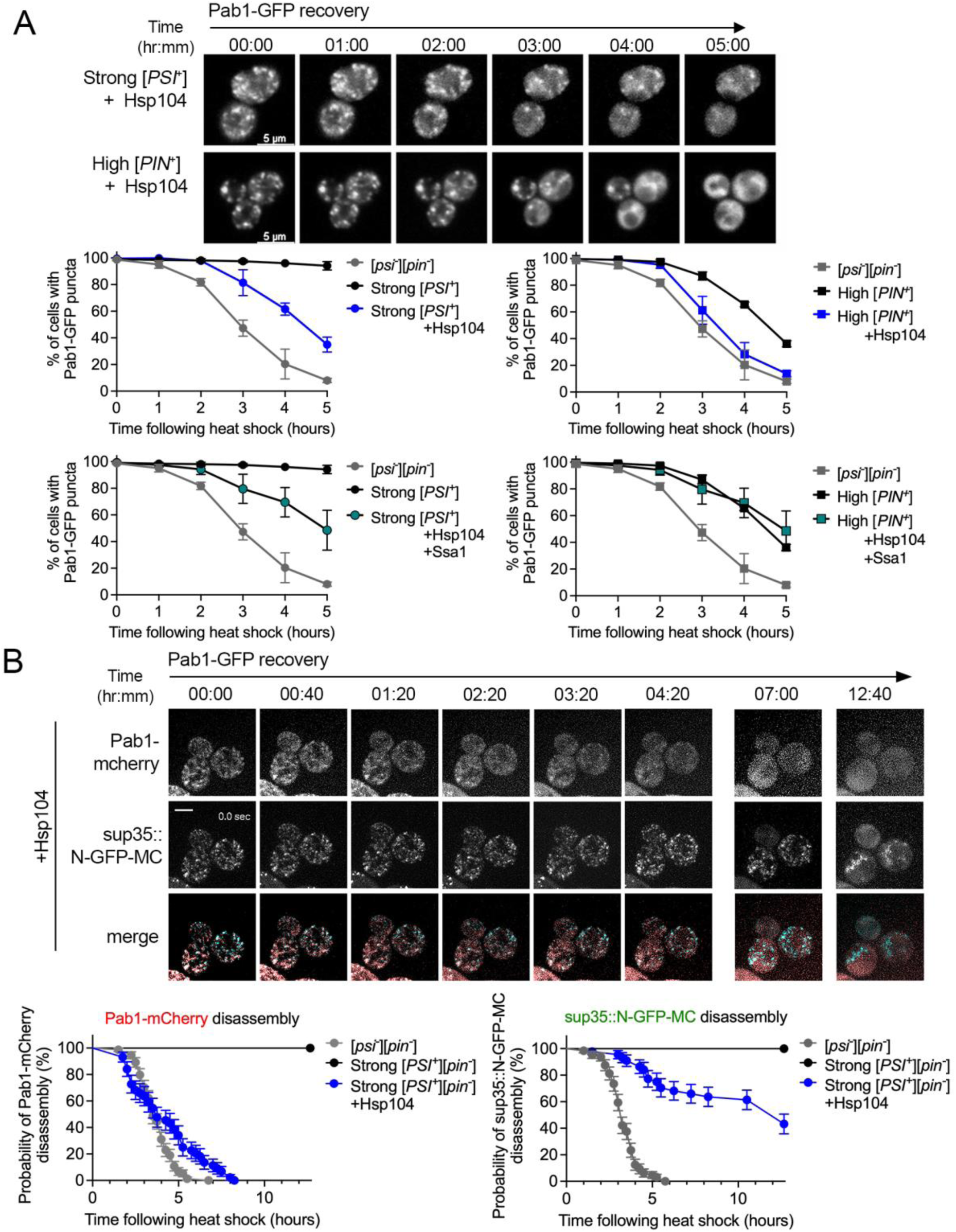
Pab1-GFP disassembly occurs in [*PSI*^+^] strains with Hsp104 overexpression. A. Top. Strong [*PSI*^+^] or high [*PIN*^+^] strains containing Pab1-GFP with a plasmid containing HSE-Hsp104 (+Hsp104) were heat shocked and subjected to 3D-timelapse microscopy. Shown are the maximum projections. Bottom. The percent of cells containing Pab1-GFP puncta were assessed from images similar to Figure 2A. Percentages were quantified from approximately 200-300 cells from 3-4 replicates at each time point (mean, ±SD). No prion ([*psi*^-^][*pin*^-^]) and prion strains from Figure 2A are included in each graph for reference. B. [*PSI*^+^] sup35::N-GFP-MC strains containing Pab1-mCherry and overexpressing Hsp104 were heat shocked and subjected to widefield 3D timelapse microscopy every 15 minutes for 12.67 hours. Shown are cells immediately following heat shock (00:00) and subsequent indicated times during recovery. Timelapse microscopy was used to quantify the time in which visible Pab1-mCherry and sup35::N-GFP-MC puncta are disassembled in no prion, strong [*PSI*^+^], and strong [*PSI*^+^] +Hsp104 strains. Stress granule disassembly was analyzed using survival curves, in which the proportion of cells retaining visible puncta is plotted over time, and the cells that have gone from containing aggregates to diffuse fluorescence as the event (n=45-75 cells per strain). Note that cells containing both Pab1-mCherry and sup35::N-GFP-MC puncta were used in this analysis. [*psi*-][*pin*-] and strong [*PSI*^+^] survival curves are included for reference.

We then tested how stress granule recovery is impacted by co-overexpression of two chaperones in strains containing prions. Unlike strains lacking prions where co-overexpression of Ssa1 and Hsp104 led to stress granule recovery similar to normal strains (Figure S8B), [*PSI*^+^] strains with co-overexpression of Hsp104 and Ssa1 resulted in disassembly kinetics similar to Hsp104 overexpression alone (Figure 4A). These data imply that in the presence of [*PSI*^+^], the competition between stress granules and prion is dependent upon Hsp104 resource availability. However, [*PIN*^+^] strains show that co-overexpression of Hsp104 and Ssa1 results in stress granule kinetics that are similar to scenarios when no chaperones are overexpressed (Figure 4A), suggesting that while Hsp104 may rescue disassembly defects in the presence of [*PIN*^+^], co-overexpression with Ssa1 interferes with this rescue.. Based on these combined results, we suspect [*PSI*^+^] and [*PIN*^+^] engage chaperone resources differently, thereby impacting how stress granules recover.

Five hours after heat shock, Hsp104 overexpression only led to partial rescue of Pab1-mCherry disassembly in [*PSI*^+^] cells (Figure 4A). Upon visualizing stress granule recovery on longer time scales with Hsp104 overexpression, the majority of cells show Pab1-mCherry diffuse fluorescence by 8.25 hours, which is only slightly slower than normal Pab1-mCherry stress granule recovery in the absence of prions (6.75 hours; Figure 4B). In these same cells, we also monitored the presence of sup35::N-GFP-MC aggregates. While most cells maintain sup35::N-GFP-MC aggregates, a subpopulation of cells displayed diffuse fluorescence with Hsp104 overexpression. We suspect that the diffuse population maintains the majority of cells in the [*PSI*^+^] state since prion plating assays show that almost all colonies are [*PSI*^+^] (Figure S7A). Since previous work suggested that non-visible particles are responsible for propagation of [*PSI*^+^]^48^, it is possible that Hsp104 overexpression may reduce the size of aggregates so that they are not visible, yet non-visible aggregates are sufficient to propagate the prion. Altogether, our data suggest that Hsp104 is associated with the prion before heat shock, thus limiting the Hsp104 available for stress granule disassembly resulting in stress granule persistence. The introduction of additional Hsp104 through overexpression is sufficient to restore stress granule disassembly in the presence of [*PSI*^+^] and [*PIN*^+^], indicating that the disassembly defect observed in prion strains possibly arises through the competition between the prion and stress granules for a shared pool of chaperone resources.

## Discussion

Using an *in vivo* model, we uncovered a resource competition mechanism that may explain why stress granule persistence occurs. In cells that do not contain amyloid or other aggregating proteins, stress granules are dynamic and disassemble upon stress release. However, there is a lack of mechanisms that explain the cellular conditions that impair disassembly and cause granules to persist. Our *in vivo* results suggest that stress granule persistence in the presence of a prion is attributed to the competition for limiting resources and can be, at least in part, restored upon chaperone addition, in this case, Hsp104. We suspect that one aggregate, such as a prion, is able to more efficiently utilize chaperone resources, thereby leaving less available for the stress granules, which may require more chaperones for disassembly. We also observed similar disassembly defects in the presence of TDP-43 aggregates, indicating that the simple presence of an initial aggregate may be sufficient to limit stress granule disassembly and foster persistence.

We propose a model based on the classical resource competition hypothesis, where two species compete for a limited resource (Figure 5). According to the hypothesis, species compete for limiting resources, and the species that make use of resources more efficiently can outcompete the other^25^. In the case of both [*PSI*^+^] and [*PIN*^+^], our data suggest that Hsp104 is the limiting resource. These prions are dependent upon the presence of Hsp104 for fragmentation since deletion of Hsp104 leads to prion loss^35, 49^. However, very high levels of Hsp104 have been shown to cure the [*PSI*^+^] prion but not [*PIN*^+^]^49, 50^. While the levels of Hsp104 in wildtype cells are sufficient to propagate prions or to separately disassemble stress granules after heat shock, it is possible that the Hsp104 chaperone requirements to carry out both fragmentation and stress granule disassembly together is beyond the amount that is available in the cell. We suspect that ongoing chaperone use through the cycling of chaperone complexes to ensure prion fragmentation may limit the availability of chaperones for stress granule disassembly. Since overexpression of Hsp104 and Ssa1 had differential effects on stress granule disassembly in [*PSI*^+^] vs. [*PIN*^+^] strains, our data suggest that each prion protein may have a different set of chaperone requirements, some of which are shared resources with stress granules that could lead to stress granule persistence.

**Figure 5.**
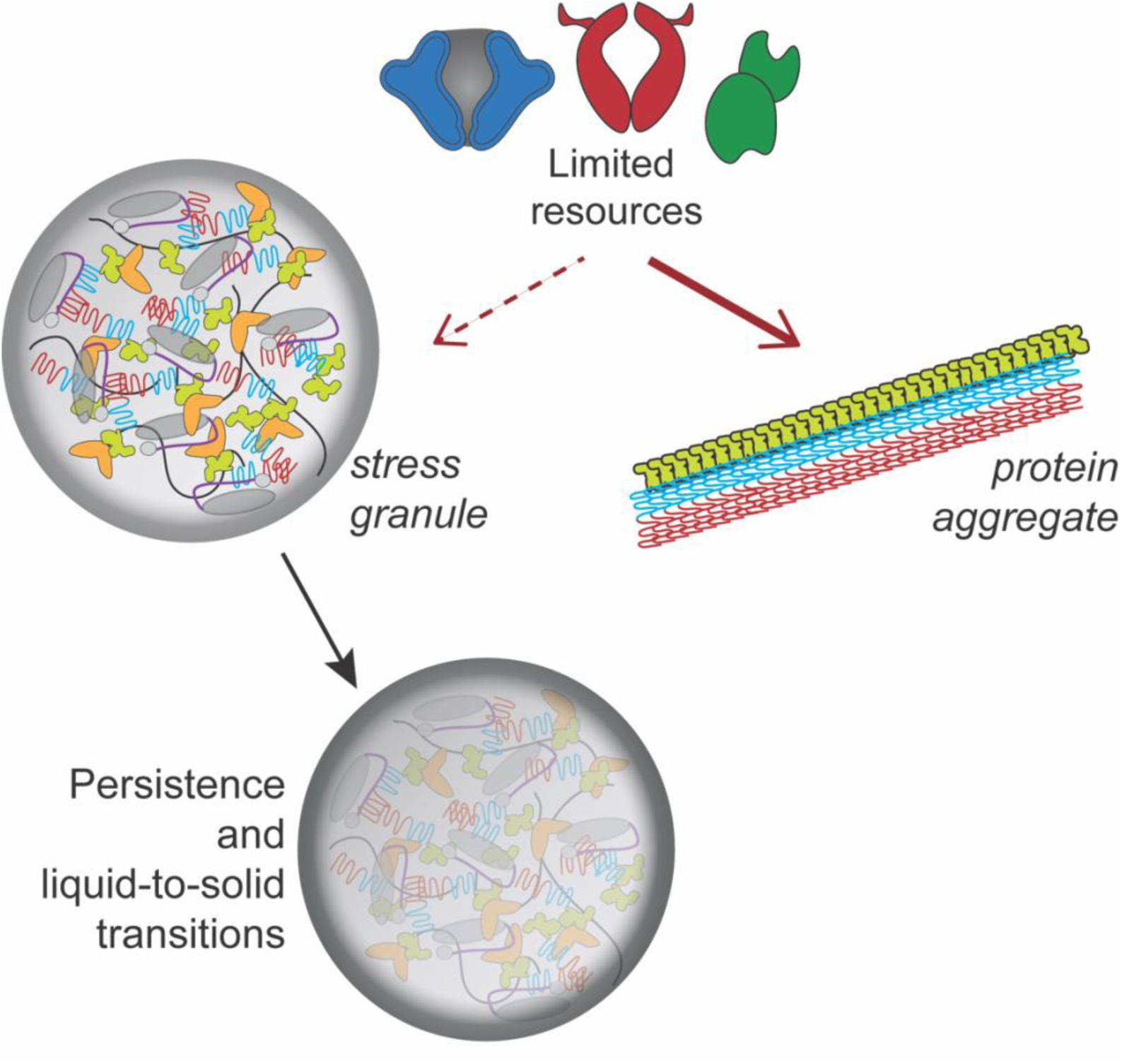
Model for stress granule persistence by resource competition. Pre-existing protein aggregates may be more efficient at utilizing limited cellular resources (as shown by solid red arrow), leaving less resources available for other cellular functions such as stress granule disassembly. The persistence of stress granules may lead to localized demixing and liquid-to-solid transitions.

Resource competition may be one of many drivers that result in stress granule persistence. In the clinic, resource competition could be critical for understanding why a portion of Alzheimer’s patients accumulate TDP-43 deposits. Our results reveal a mechanism that explains why stress granule disassembly can be impaired resulting in further stress granule persistence leading to solidification and further aggregation. Future studies are warranted to identify and understand the unique resources shared between stress granules and aggregating proteins, in order to provide new directions for addressing neurodegenerative disease states.

## Methods

### Strains and cultivation procedures

*S. cerevisiae* strains utilized in this study were in the 74D-694^49^ or BY4741 genetic background^51^. Cells were propagated in either yeast peptone dextrose (YPD) media or synthetic complete (0.17% yeast nitrogen base and 0.5% ammonium sulfate) media lacking the indicated amino acids and containing either 2% dextrose, 2% raffinose, or 2% raffinose and a specified percentage of galactose. Strains were grown at 30°C overnight in round bottom 50 ml Oak Ridge tubes using plasmid-selective liquid media or in 250ml flasks with shaking. Overnight growth ranged from 16-18 hours until late log phase (OD_600_ 0.8-1.2). For galactose inductions, strains were grown overnight in SD media, washed three times in sterile water, and then inoculated at OD_600_ 0.1 in 2% raffinose or 2% raffinose/galactose media until OD_600_ 0.5-0.8. Heat shock was conducted by transferring 300μL of overnight cultures into a 1.5 mL microfuge tube, incubated at 42°C for 45 minutes (or other temperature as specified) in a dry heating block for acute heat shock, or held at room temperature (no heat shock, approximately 22-25°C). Following heat shock treatments, microfuge tubes were returned to room temperature for measurements at specific timepoints or timelapse microscopy. Strains and plasmids used in this study are found in Tables S3 and S4, respectively.

### Microscopy

#### Widefield fluorescent microscopy

To quantify SG disassembly via snapshots of fields of cells during recovery, indicated strains containing either a Pab1-GFP or Pab1-mCherry plasmid were grown overnight to late log phase at 30°C. Cells were heat shocked as described above. Images were taken of cells immediately following heat shock and at each hour during the five hours of recovery at room temperature. Images were acquired with approximately 13 Z-stacks with Leica DMI 6000 fluorescent deconvolution microscope (630X, 1.4 NA) and captured with a Leica K5 camera. The percent of cells with puncta and/or foci was determined by scoring the number of cells with visible aggregates compared to the total number of cells with fluorescence.

#### 3D-timelapse microscopy

To follow stress granule recovery over time, cells were heat shocked and 200 μL of culture was immediately added to a concanavalin A-coated Ibidi 8-well glass bottom slide. For two-color microscopy, images were captured similar to above using the Leica DMI 6000 microscope. For three-color microscopy, images were captured using a Zeiss AxioObserver (100X, 1.44 NA) with SpectraX light source and captured with ORCA Flash CMOS camera using VisiView Software with Visiboost.

#### Spinning disk confocal microscopy

To detect tagged Pab1, Sup35, and Hsp104 puncta and/or foci, indicated strains were imaged with a Zeiss AxioObserver, CrestOptics Cicero spinning-disk confocal system (100X, 1.44 NA), and captured with ORCA Flash CMOS camera. Images were processed using VisiView software (Visitron Systems GmbH, Puchheim, Germany). Snapshot images were assessed by analyzing the indicated number of cells that contained either diffuse GFP fluorescence or exhibited fluorescent puncta. Pearson’s Correlation Coefficients and line plots were derived from overlays of the indicated channels from the middle Z-stack of indicated 3D images using ImagePro 11 software.

#### Near-TIRF microscopy

For imaging of Pab1-mCherry and Sup35-GFP at reduced signal-to-noise ratio, indicated strains were inoculated in SD medium lacking tryptophan and supplemented with 0.04 mg/ml adenine and grown at 30°C in a shaking incubator until they reached an OD_600_ of approximately 0.7. Subsequently, cells were pelleted and resuspended in 50 µl of medium. Paired cultures were either incubated at room temperature or heat shocked at 42°C for 45 min. Images were collected in near-TIRF mode ^52^ with a Leica DMi8 inverted fluorescence microscope equipped with a 100X, 1.47 NA Plan-Apochromat oil immersion objective, a 1.6X magnification changer, a Flash 4.0 v3 sCMOS camera (Hamamatsu), a W-View Gemini beam splitter (Hamamatsu), 488 and 561 nm lasers, a Quad-T excitation/emission filter, and LAS X v3.7.6.25997 software.

### Fluorescence Recovery after Photobleaching

Paired cultures were grown as described and either left untreated at room temperature or heat shocked for 45 minutes at 42°C. Heat shocked samples were imaged immediately after heat shock. For recovery samples, cultures were allowed to incubate at room temperature for 2.5 hours until imaged. Cells were imaged on a Nikon A1R inverted Eclipse Ti confocal microscope using a Plan-Apochromat 60x water objective. The pinhole was set for 14.0μm/0.4 airy units on the 488nm laser. Images were acquired pre-bleach, and the ROIs were photobleached using a 488nm laser at 100% for 2.1 seconds. Fluorescence recovery was monitored for 3 minutes at 2.1 seconds/frame after bleaching. Image analysis was conducted using Image Pro 11. ROIs were taken from 1) the region of bleaching, 2) the region within the same cell that was not bleached, and 3) outside the cell. The corrected fraction intensity was determined according to Wu et al.^53^. Curves were drawn using a non-linear regression exponential curve with two phase separation using GraphPad Prism V. 10.6.1. A minimum of seven trials were acquired for each treatment (Table S2).

### Quantification and statistical analysis

Graphs were generated using GraphPad Prism V. 10.6.1. Statistical analysis was performed using Prism and statistical tests are indicated in the figure legends. To determine Pearson’s correlation coefficient (PCC), a 150×150 pixel area was cropped and both channels were merged in ImagePro 11. PCC values were derived from the middle Z-stack using ImagePro 11 software. A minimum of 30 cells were analyzed per strain.

## Supporting information

Supplemental Figures, Tables, and Raw data

## Acknowledgements

Special thanks to Stefan Schnitzer (Smithsonian TRI/Marquette) for fruitful discussions about resource competition in ecology, Lisa Petrella for advice to strengthen our quantitative analysis, and Claire Radtke for critical review of the manuscript. The authors would like to thank Roy Parker (UC Boulder), Susan Liebman (UN-Reno), and David Pincus (U Chicago) for plasmids, Susan Liebman (UN Reno) and Tricia Serio (UWash) for strains, and Elizabeth Craig (UW-Madison) for antibodies used in these studies. The BE4 (Sup35C) antibody was a gift from Viravan Prapapanich and Susan Liebman. This work was supported by the National Science Foundation (MCB 2127616) and the National Institutes of Health (GM155860) to ALM, National Science Foundation (MCB CAREER 1942395) to DCP, and Marquette University Way Klingler Startup funds to EMS. HEB was supported by the Marquette Raynor Fellowship.

## Notes

### Competing Interest Statement

The authors have declared no competing interest.

