## Supplemental Figures, Tables, and Raw data for "Stress granules and protein aggregates reveal intracellular resource competition"

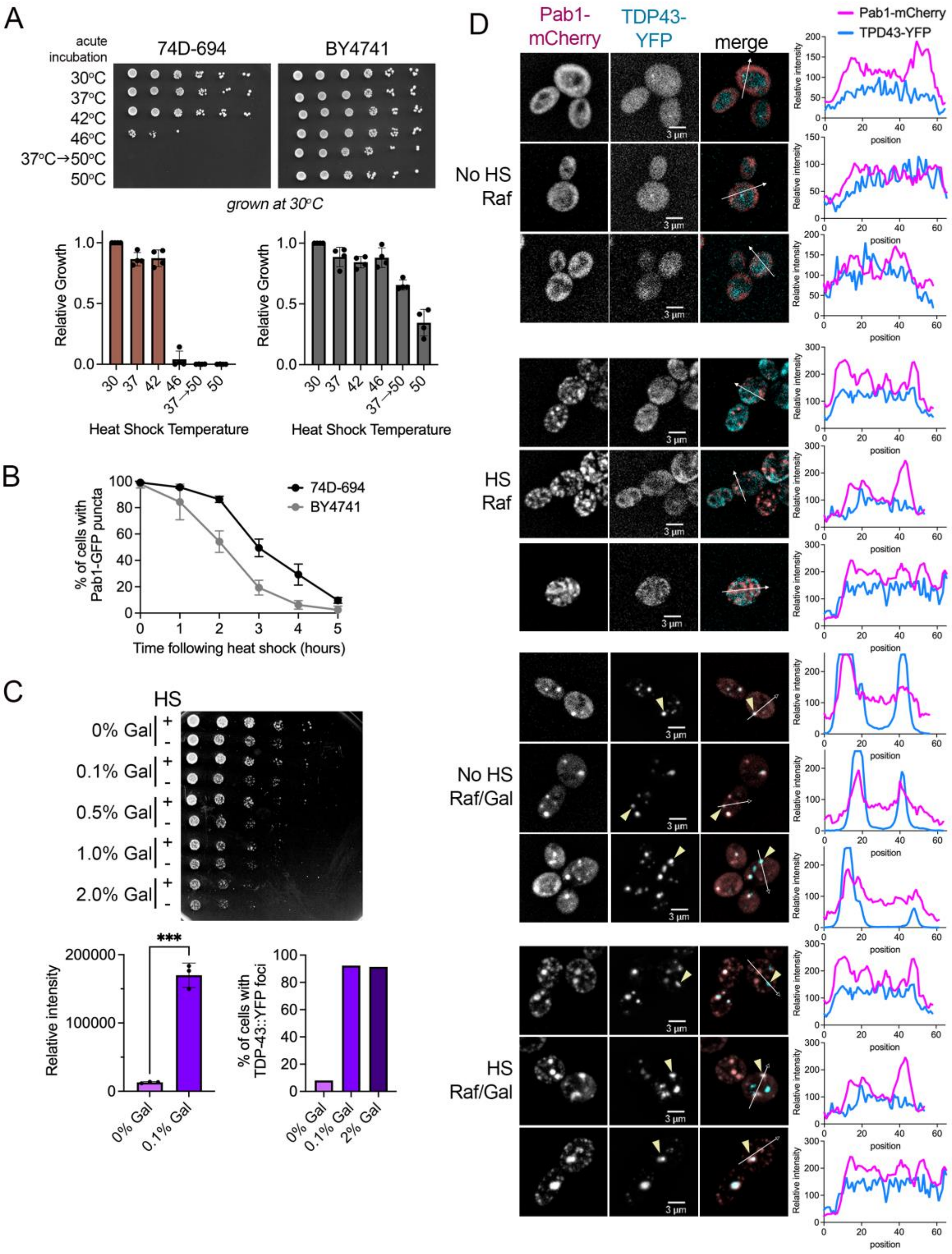

A
acute incubation
74D-694
BY4741
30°C
37°C
42°C
46°C
37°C→50°C
50°C
grown at 30°C
Relative Growth
Heat Shock Temperature
B
% of cells with Pab1-GFP puncta
74D-694
BY4741
Time following heat shock (hours)
C
HS
0% Gal
0.1% Gal
0.5% Gal
1.0% Gal
2.0% Gal
Relative intensity
***
% of cells with TDP-43::YFP foci
2% Gal
D
Pab1-mCherry
TDP43-YFP
merge
Pab1-mCherry
TPD43-YFP
No HS Raf
HS Raf
No HS Raf/Gal
HS Raf/Gal
3 μm
Relative intensity
position

**Figure S1. Heat stress response varies in different genetic backgrounds, and induction of TDP-43 aggregation.** A. *Top*. 74D-694 or BY4741 strains were incubated at the indicated temperatures for 45 minutes (acute incubation) and serially diluted five-fold and plated. Thermotolerance assays (37°C→50°C) were conducted by incubating strains at 37°C for 30 minutes followed by 50°C for 30 minutes before plating [1]. *Bottom*. Graphs show quantification of relative growth for four biological replicates with mean ±SD for 74D-694 (left) and BY4741 (right) [2]. B. 74D-694 or BY4741 strains were heat shocked at 42°C for 45 minutes and allowed to recover at room temperature. Images were taken of populations at the indicated times. The percentage of cells with Pab1-GFP stress granules after heat shock treatment was determined for each time point (200-300 cells per replicate, n=3, mean ±SD). C. Top. [*psi*-][*pin*-] strains containing Pab1-mCherry and a galactose inducible TDP-43-YFP were grown in media overnight containing 2% raffinose and the indicated percentages of galactose (Gal). Cultures were then either incubated at room temperature (-) or heat shocked at 42°C for 45 minutes (+), serially diluted five-fold and plated on selective media containing 2% dextrose. Bottom left. Three independent cultures containing TDP-43-YFP were grown in 2% raffinose (0% Gal) or 2% raffinose/0.1% galactose (0.1% Gal) and assessed for YFP intensity through flow cytometry using the FITC filter. Mean FITC area of three individual trials were compared by an unpaired t-test (***$p<0.001$). Bottom right. No heat shock cultures in 2% raffinose and the indicated percentage of galactose were visualized by fluorescent microscopy and the percentage of cells containing TDP-43-YFP foci was quantified. D. Strains grown without galactose (Raf) or with 0.1% raffinose (Raf/Gal) were incubated at room temperature (No HS) or heat shocked (HS) conditions. Spinning disk confocal microscopy was used to image cells. Line plots graph the relative fluorescent intensity along the line drawn in the merged image. Examples of large TDP-43-YFP aggregates are indicated by yellow arrows.

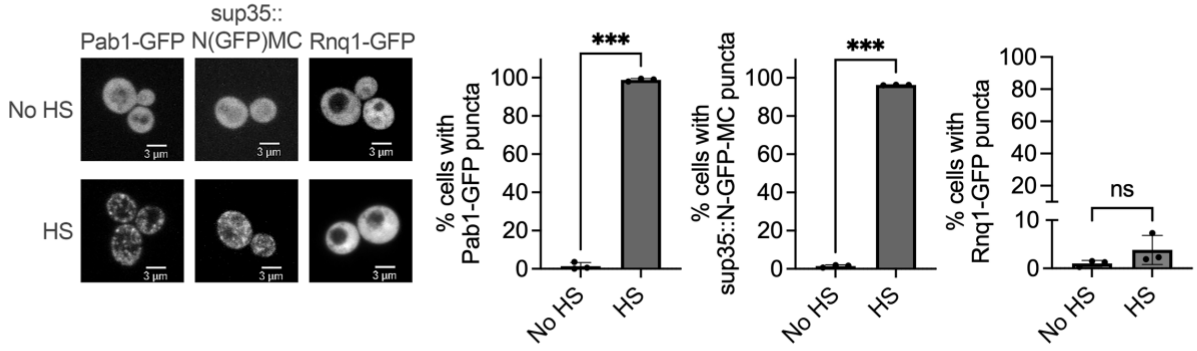


**Figure S2. Sup35 is heat responsive, but Rnq1 is not.** 74D-694 [*psi*-][*pin*-] strains expressing Pab1-GFP, sup35::N-GFP-MC or Rnq1-GFP were imaged under no heat shock conditions (No HS) or immediately after a heat shock of 42$^{o}$C for 45-minutes (HS). The percentage of cells with puncta were quantified. Comparisons between treatments or strains were compared by an unpaired t-test with Welch's correction (***p<0.001). Each dot represents the mean of 200-300 cells per replicate (n=3, mean ±SD).

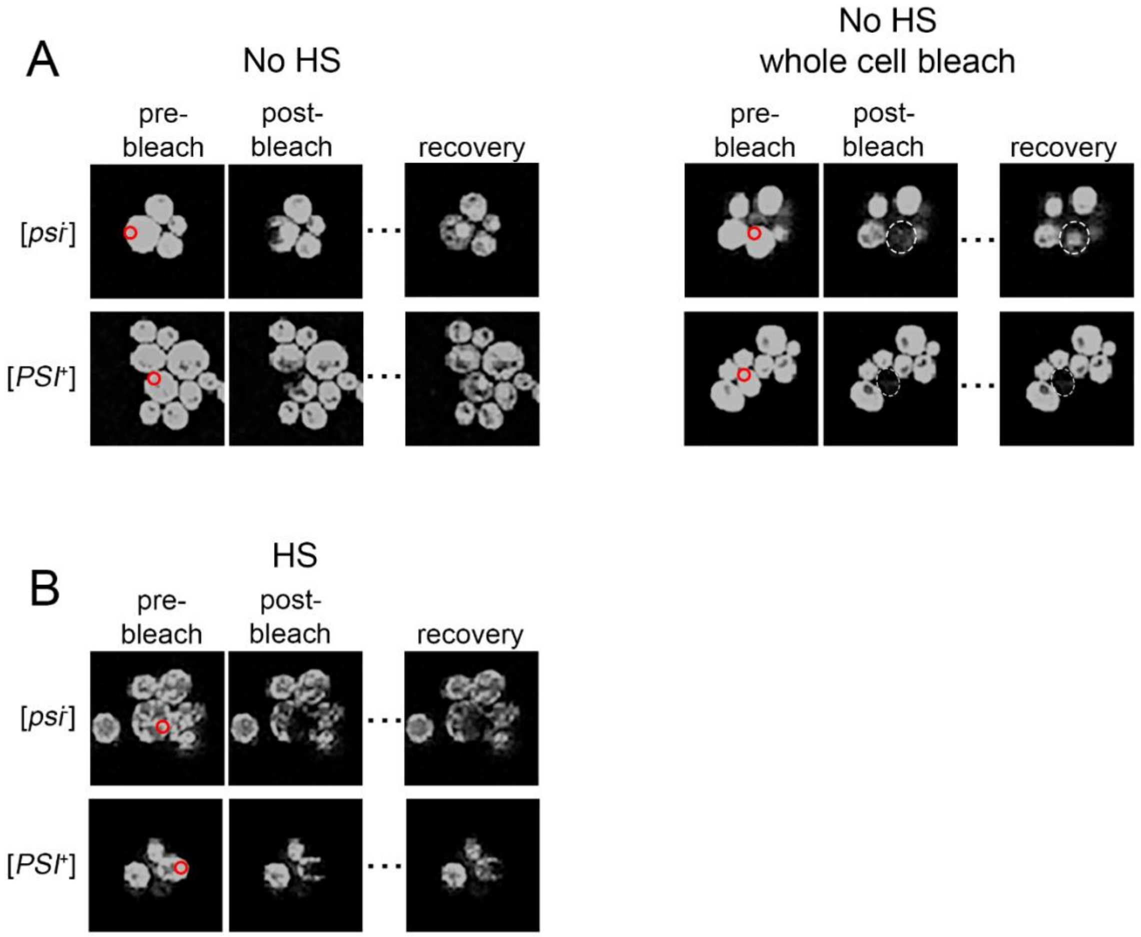


**Figure S3. Images of cells during FRAP.** A. Images depicting FRAP analysis of Pab1-GFP in [$psi^-$] or strong [$PSI^+$] strains that were untreated (no HS). Images show fluorescence before (pre-bleach), immediately after bleaching (post-bleach) and 342 seconds after photobleaching (recovery). Representative images are shown. Red circle indicates the region targeted for bleaching. Note that a proportion of non-heat shocked cells displayed complete cellular bleaching immediately after bleaching Table S2). The white dotted circles indicate the cells that were fully bleached. B. FRAP conducted on [$psi^-$] or strong [$PSI^+$] strains immediately after heat shock (HS).

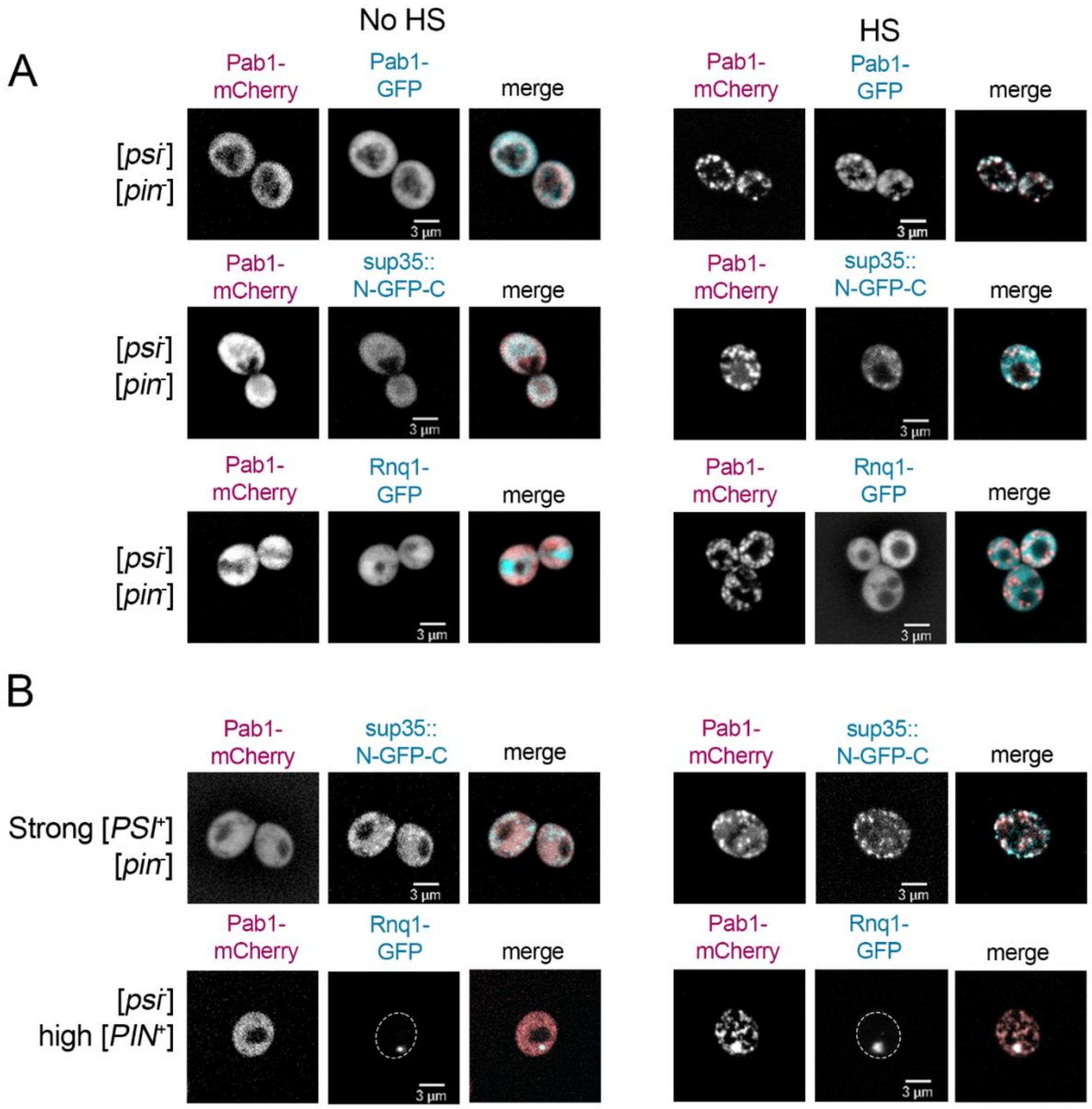


**Figure S4. Colocalization images of Pab1 and Sup35 or Rnq1 under no heat shock or after heat shock.** A. Top. No prion ([*psi*$^-$][*pin*$^-$]) cells expressing Pab1-GFP and Pab1-mCherry were imaged by spinning disk confocal microscopy under no heat shock conditions (No HS, left) or immediately after heat shock (HS, right), and used for Pearson's correlation coefficient (PCC) control values in Figure 1A and B. No prion strains containing Pab1-mCherry, and sup35::N-GFP-MC (middle) or Rnq1-GFP (bottom) were imaged and are representative of images used in PCC analysis in Figure 1 A and B. B. [*PSI*$^+$] strains (strong variant) expressing sup35::N-GFP-MC (top) or [*PIN*$^+$] strains (high variant) expressing Rnq1-GFP (bottom) were imaged similar to A. Images were acquired with 630X magnification and shown as maximum projections.

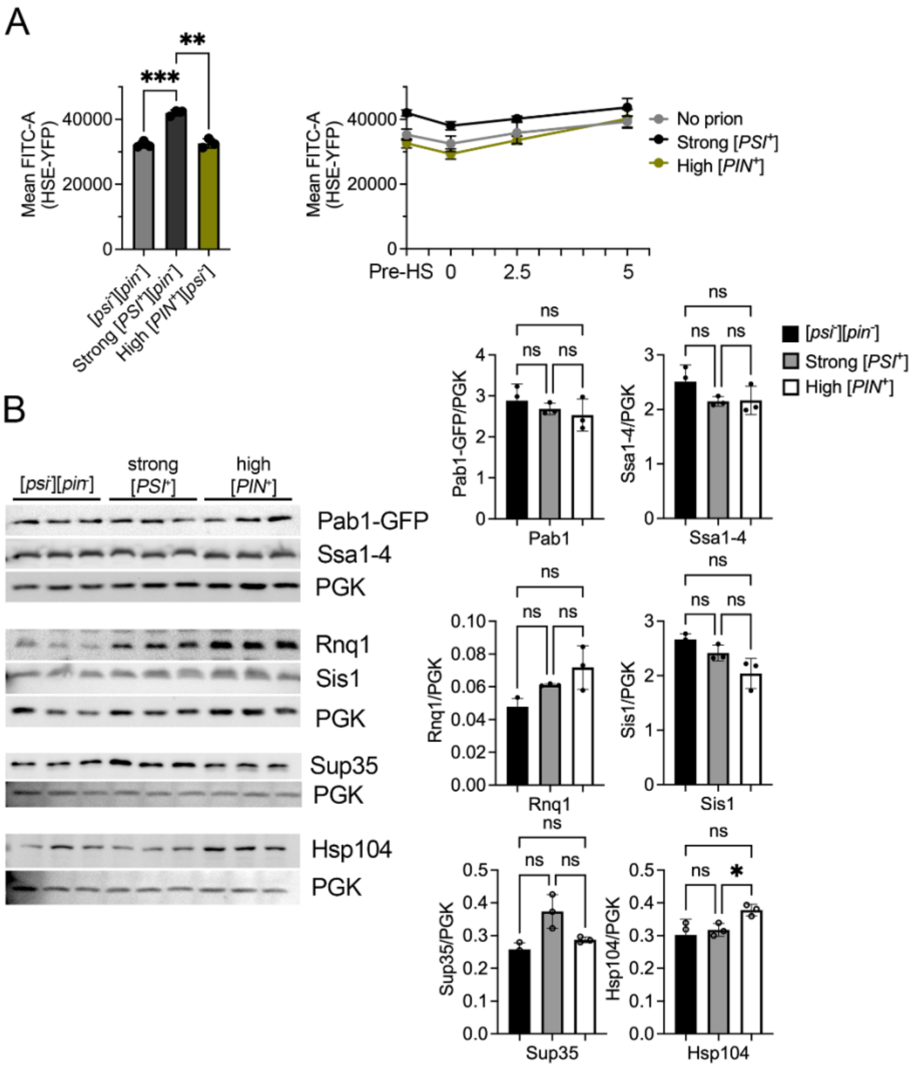


**Figure S5. [*PSI*+] strains have elevated heat shock response and Hsp104 steady state levels.** A. *Left.* An integrated HSE-YFP heat shock response reporter [3] was integrated into the indicated strains. Strains were grown to late log and assessed for HSE-YFP intensity by flow cytometry using the FITC filter in the absence of heat shock. Samples were compared using a one-way ANOVA with Games-Howell's multiple comparisons test (n=3, $^{**}p<0.01$, $^{***}p<0.001$). *Right*. Strains were then heat shocked at 42°C for 45 minutes and then allowed to recover at room temperature. Strains were monitored for FITC intensity before (pre-HS), immediately after HS (zero), and 2.5 hours and 5 hours after heat shock release. B. Fresh lysates from three independent cultures of each strain were subjected to Western blot analysis and assessed for steady-state levels of the indicated proteins when normalized to PGK levels. One-way ANOVA with Brown-Forsythe post-hoc analysis was used to compare steady state levels ($^{*}p<0.05$).

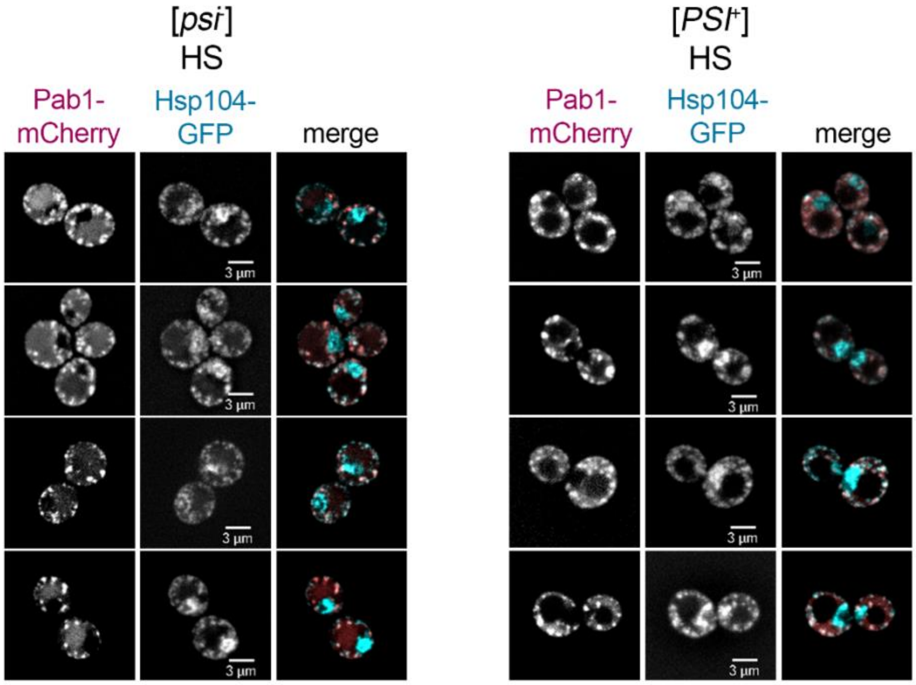


**Figure S6. Pab1 and Hsp104 colocalize in [*psi*⁻] and [*PSI*⁺] cells immediately after heat shock.** No prion ([*psi*⁻][*pin*⁻], left) or prion (strong [*PSI*⁺], right) strains containing an integrated Hsp104-GFP and transformed with Pab1-mCherry were imaged by confocal spinning disk microscopy immediately after heat shock. Images acquired with 630X magnification and shown as a maximum projection image.

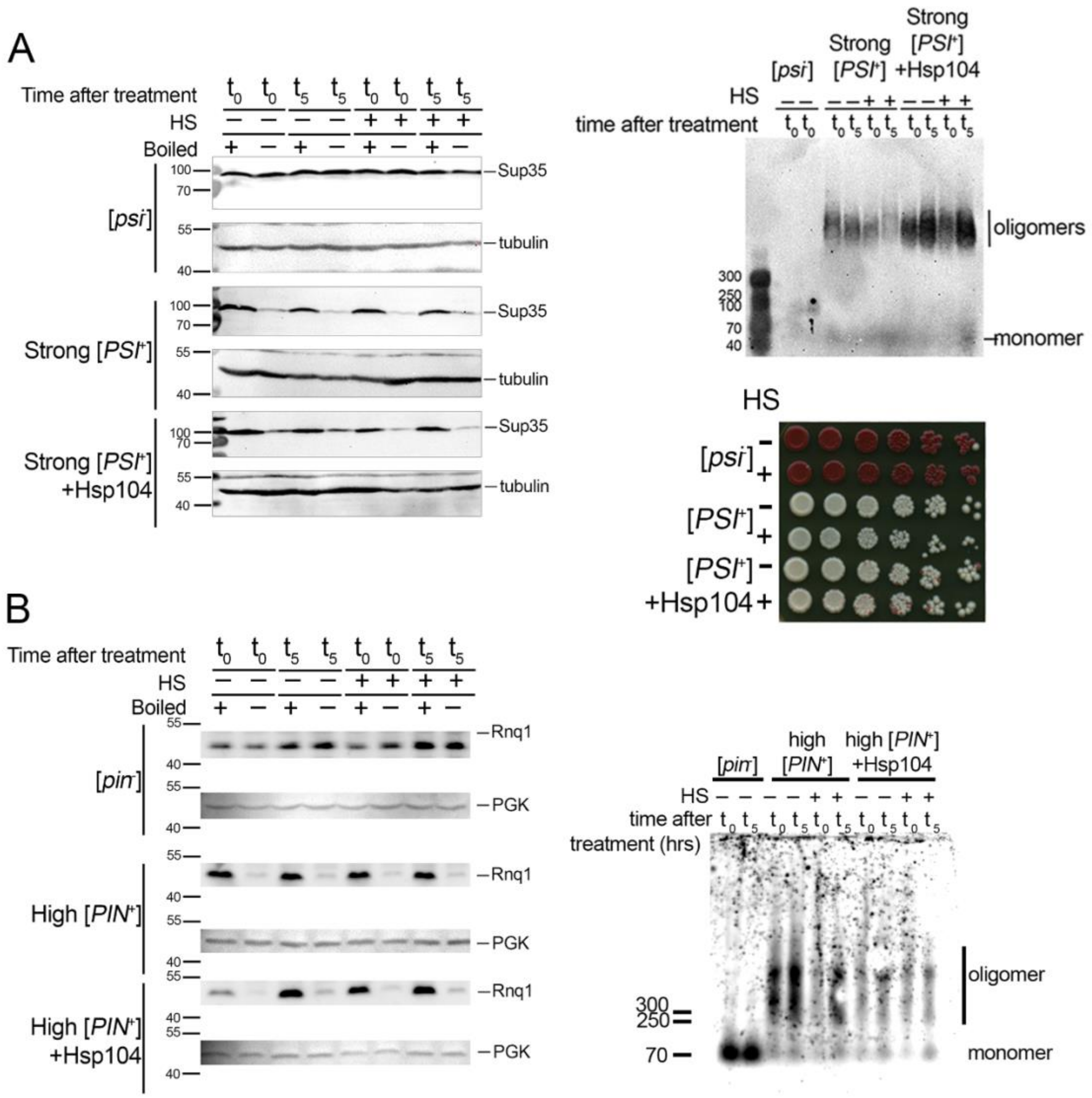


**Figure S7. [*PSI*$^+$] or [*PIN*$^+$] is maintained immediately after and 5 hours after heat shock.** A. Well trap (left) and SDD-AGE (right, top) assays of [*PSI*$^+$] strains were performed on cultures with and without Hsp104 overexpression. Strains were untreated (-) or heat shocked (+), and lysates were either taken immediately after room temperature or heat shock incubation ($t_0$) or allowed to recovery for 5 hours ($t_5$). For well-trap assays, lysates were either incubated at room temperature (-) or boiled at 98°C (+) for 8 minutes prior to loading. Images are representative of three trials. For SDD-AGE assays, lysates were not boiled. Sizes of oligomeric and monomeric protein are indicated. For [*PSI*$^+$] color assays (right, bottom), strains were either untreated (-) or subjected to heat shock (+). Cultures were serially diluted five-fold, plated on rich media, and were allowed for color to develop for 4-6 days after colonies appeared. D. Same as A but for [*PIN*$^+$] strains.

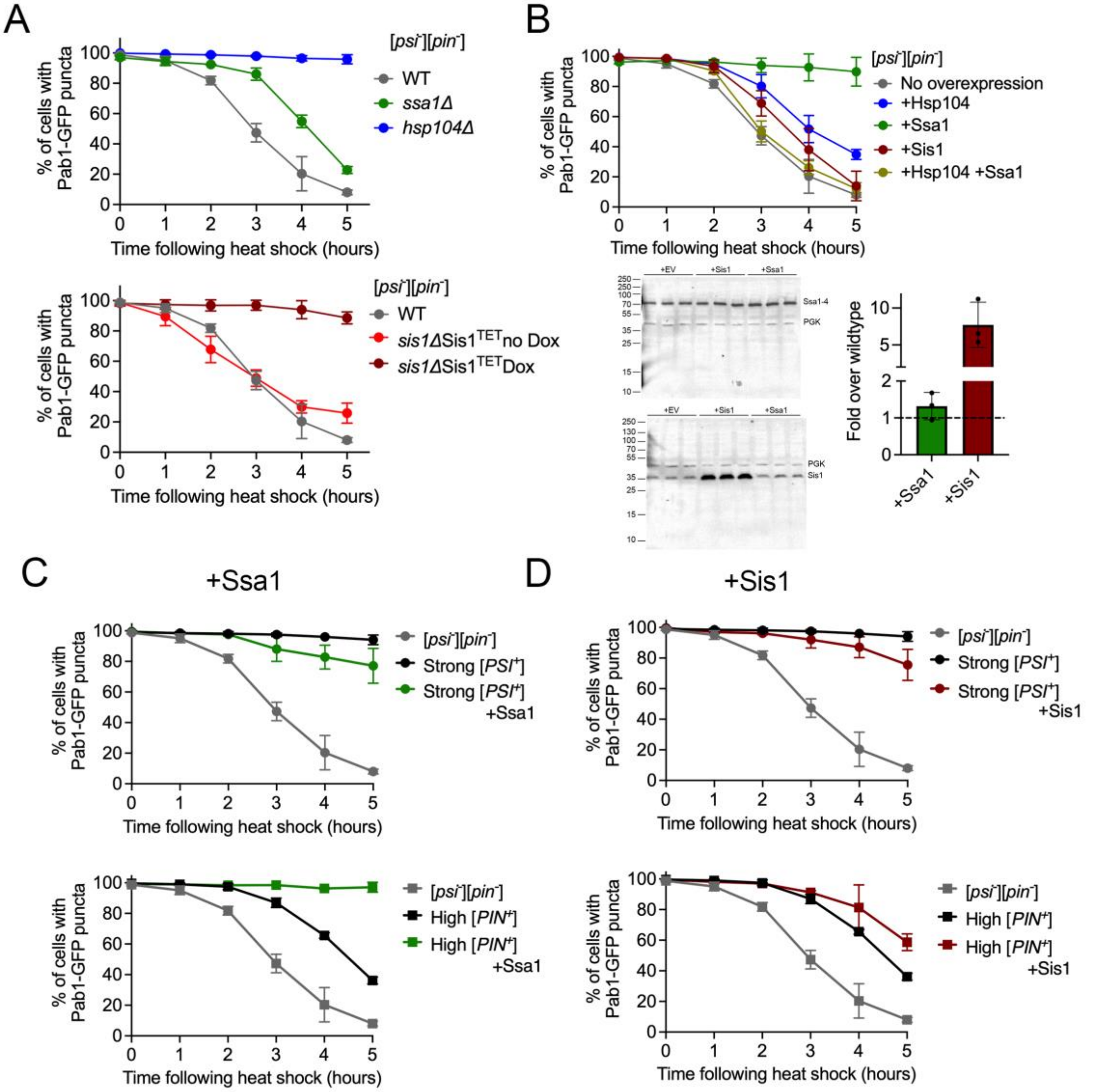


**Figure S8. Stress granule disassembly kinetics in the presence or absence of molecular chaperones.** A. The percentage of cells with Pab1-GFP puncta were monitored over time after heat shock in strains lacking *hsp104Δ* or *ssa1Δ* (top) or *SIS1* depletion where *sis1Δ* strains contain a plasmid in which Sis1 expression is controlled by a tetracycline-repressible promoter (bottom). The presence of 10mM doxycycline (Dox) represses *SIS1* expression. Stress granule disassembly was monitored similar to figure 2A (n=3-4; mean, ±SD). The no prion ([*psi*-][*pin*-]) and prion control data from Figure 2A is reproduced in each graph for consistent comparison. B. HSE-Hsp104, GPD-Ssa1, GPD-Sis1, or both HSE-Hsp104 and GPD-Ssa1 were introduced into [*psi*-] strains and monitored for Pab1-GFP puncta over time after heat shock. HSE-Hsp104 provides four-fold steady state levels over wildtype, [4] Steady state levels for GPD-Ssa1 and GPD-Sis1 was determined by Western blot quantification from three independent strains. Ssa1 or Sis1 levels were normalized over PGK, n=3. C. Pab1-GFP stress granule recovery in the indicated strains with and without GPD-Ssa1 overexpression. D. Same as C except with GPD-Sis1 overexpression.

| Strain | FRAP sample | Mobile fraction | Immobile fraction | $t_{1/2}$ (s) |
|---|---|---|---|---|
| [*psi*$^-$] | No HS | 66.7% | 33.3% | 41.9 |
| [*psi*$^-$] | Immediately after HS | 12.7% | 87.3% | 119.1 |
| [*psi*$^-$] | 2.5 hours after HS | 53.1% | 46.9% | 59.5 |
| [*PSI*$^+$] | No HS | 67.3% | 32.7% | 35.0 |
| [*PSI*$^+$] | Immediately after HS | 27.9% | 72.1% | 153.9 |
| [*PSI*$^+$] | 2.5 hours after HS | 22.3% | 77.7% | 39.0* |

**Table S1. Immobile fractions and half-life of Pab1-GFP from FRAP analysis.** Non-linear regression analysis from data in Figure 2B was used to determine the best-fit plateau of each line. The percentage below the plateau determines the proportion of the Pab1-GFP population that is mobile (M), and the percentage above the plateau is the proportion that is immobile (1-M). The half-time of recovery ($t_{1/2}$) is the time required for the bleached area to recover by 50%. *Since almost no fluorescence recovery is observed in [*PSI*$^+$], HS 2.5-hour recovery cells, $t_{1/2}$ values are irrelevant.

| Strain | FRAP samples | Total number of cells analyzed | Number of cells that showed complete bleaching |
|---|---|---|---|
| [*psi*$^-$] | No HS | 23 | 14 |
| [*psi*$^-$] | Immediately after HS | 19 | 0 |
| [*psi*$^-$] | 2.5 hours after HS | 7 | 1 |
| [*PSI*$^+$] | No HS | 19 | 11 |
| [*PSI*$^+$] | Immediately after HS | 17 | 1 |
| [*PSI*$^+$] | 2,5 hours after HS | 8 | 1 |

**Table S2. A proportion of no heat shocked cells resulted in complete bleaching of the cell.** The total number of cells analyzed by FRAP analysis. Under no heat shock conditions, a large proportion of cells that were photobleached in a region of interest resulted in complete absence of GFP signal from the entire cell (as shown in Figure S3A). The complete bleaching suggests that almost all of the mobile Pab1-GFP cytosolic protein is turned over despite the bleached area being only a small portion of the cell.

| Strain # | Genotype | Name | Reference |
|---|---|---|---|
| D230 | *Mat**a** ade1-14 ura3-52 leu2-3,112 trp1-289 his3-200* [*psi*-][*pin*-] | [*psi*-][*pin*-] 74D-694 | [5] |
| M101 | *Mat**a** leu2Δ his3Δ ura3Δ met15Δ* [*psi*-] [*pin*-] | [*psi*-][*pin*-] BY4741 | [6] |
| D114 | *Mat**a** ade1-14 ura3-52 leu2-3,112 trp1-289 his3-200* Strong [*PSI*+][*pin*-] | Strong [*PSI*+] 74D-694 | [7] |
| D233 | *Mat**a** ade1-14 ura3-52 leu2-3,112 trp1-289 his3-200* [*psi*-] High [*PIN*+] | High [*PIN*+] 74D-694 | [8] |
| M703 | *Mat**a** ade1-14 ura3-52 leu2-3,112 trp1-289 his3-200* [*psi*-][*pin*-] 4xHSE-YFP::LEU2 | [*psi*-][*pin*-] 74D-694 HSE-YFP | [9] |
| M704 | *Mat**a** ade1-14 ura3-52 leu2-3,112 trp1-289 his3-200* high [*PIN*+][*psi*-] 4xHSE-YFP::LEU2 | High [*PIN*+] 74D-694 HSE-YFP | This study |
| M706 | *Mat**a** ade1-14 ura3-52 leu2-3,112 trp1-289 his3-200* strong [*PSI*+][*pin*-] 4xHSE-YFP::LEU2 | Strong [*PSI*+] 74D-694 HSE-YFP | This study |
| M716 | *Mat**alpha** ade1-14 UGA/ trp1-289 his3-del200 ura3-52 leu2-3,112*/ sup35::N(GF) 3sGFP (GS)3MC [*pin*-] [*psi*-] | [*psi*-][*pin*-] 74D-694 Sup35::N-GFP-MC | This study |
| M719 | *Mat**alpha** ade1-14 UGA/ trp1-289 his3-del200 ura3-52 leu2-3,112*/ sup35::N(GF) 3sGFP (GS)3MC strong [*PSI*+][*pin*-] | Strong [*PSI*+] 74D-694 Sup35::N-GFP-MC | This study |
| M758 | *Mat**alpha** ade1-14 UGA/ trp1-289 his3-del200 ura3-52 leu2-3,112*/ sup35::N(GF) 3sGFP (GS)3MC [*pin*-][*psi*-] Hsp104-mTagBFP2::LEU2 | [*psi*-][*pin*-] 74D-694 Sup35::N-GFP-MC Hsp104-BFP | This study |
| M760 | *Mat**alpha** ade1-14 UGA/ trp1-289 his3-del200 ura3-52 leu2-3,112*/ sup35::N(GF) 3sGFP (GS)3MC strong [*PSI*+][*pin*-] Hsp104-mTagBFP2::LEU2 | Strong [*PSI*+] 74D-694 Sup35::N-GFP-MC Hsp104-BFP | This study |

**Table S3. Yeast Strains used in this study**

| Plasmid # | Description | Yeast marker | Reference |
|---|---|---|---|
| p3284 | Pab1-GFP | *URA3* | [10] |
| p3285 | Pab1-GFP | *TRP1* | [10] |
| p3110 | HSE-Hsp104 | *URA3* | [11] |
| p3169 | HSE-Hsp104 | *LEU2* | [12] |
| p3302 | GPD-Ssa1 | *LEU2* | This study |
| p3281 | GPD-Sis1 | *TRP1* | This study |
| p3387 | HSE-YFP | *LEU2* | [3] |
| p3384 | Pab1-mCherry | *TRP1* | This study |
| p3036 | Rnq1-GFP | *LEU2* | [13] |
| p3386 | Hsp104-mTagBFP2 | *LEU2* | [14] |
| p3071 | GAL-TDP-43::YFP | *URA3* | [15] |
| p3011 | GAL-YFP | *URA3* | [16] |

**Table S4. Plasmids used in this study.**

| Antibody | Dilution | Clonality | Vendor |
|---|---|---|---|
| Anti-GFP | 1:5000 | Monoclonal | Roche (clones 7.1 and 13.1) |
| Anti-Sup35C | 1:10000 | Monoclonal | BE4 antibody was a kind gift from Viravan Prapapanich and Susan Liebman (UN-Reno) |
| Anti-Hsp104 | 1:10000 | Polyclonal | Enzo (ADI-SPA-1040) |
| Anti-Rnq1 | 1:1000 | Polyclonal | Elizabeth Craig (UW-Madison) |
| Anti-Ssa1-4 | 1:10000 | Polyclonal | Elizabeth Craig (UW-Madison) |
| Anti-Sis1 | 1:10000 | Polyclonal | Elizabeth Craig (UW-Madison) |
| Anti-PGK | 1:10000 | Monoclonal | Abcam (22C5D8) |

**Table S5. Antibodies used in this study**

### Supplemental Materials and Methods

*Growth assays*

Cultures were grown in liquid media at 30$^o$C overnight to late log phase. Cultures were normalized to an $OD_{600}$ of 0.6 and incubated at the indicated temperatures for 45 minutes. Thermotolerance assays were performed as described in Sanchez and Lindquist [1], where cultures were incubated at 37°C for 30 min, followed by an additional heat treatment at 50°C for 30 minutes. Cultures were serially diluted five-fold and spotted on SD-Ura media. The plates were incubated at 30°C for 2-3 days. Quantification of relative growth was performed according to Petropavlovskiy et al.. [2].

*Flow cytometry*

Strains containing Pab1-GFP or HSE-YFP were grown overnight in liquid media to late log. Flow cytometry was performed using a Cytoflex Flow Cytometer (Beckman Coulter) to measure GFP fluorescence intensity (FITC filter). 50,000-100,000 cells were counted per sample. Histograms and scatterplots and statistical means were generated using CytExpert Software.

*FRAP analysis*

The mobile, immobile and time-to-half-recovery was based on Sprunger and Jackrel[17]. Briefly, a line is drawn the plateau of the nonlinear regression curve. The calculated value under the plateau is the mobile fraction, and the calculated value above the plateau is the immobile fraction. The time-to-half recovery is the time required for 50% of the fluorescent signal to recover to half of the final fluorescent intensity.

*Western Blot Analysis*

Strains were grown overnight to late log phase at 30°C. Lysates were prepared by cold glass bead lysis as previously described[18]. 25μg of lysate was treated with 2% SDS sample buffer (25 mM Tris, 200 mM glycine, 5 % glycerol, and 0.025 % bromophenol blue) and heated to 95°C for 7 minutes. Proteins were resolved on 10% SDS PAGE and blots were subjected to normal Western blotting procedures. Antibodies used in this study are found in Table S5. Images were captured with the iBright 1500 (ThermoFisher) and quantified in ImageJ or iBright software. All signals were normalized to loading controls.

*Prion propagation detection*

*Well trap assays.* 25 μg of lysates was diluted in 1X in 2% SDS sample buffer. Samples were either incubated at room temperature (unboiled) or 95°C for 10 minutes (boiled) and loaded onto a 10% SDS-PAGE. Blots were subjected to Western blot analysis.

*SDD-AGE.* 100 μg of prepared lysates was prepared in 1X lysis buffer and incubated at room temperature for 10 minutes. Samples were run on 1.5% agarose gel and transferred to PVDF according to Halfmann and Lindquist [19]. Blots were subjected to Western blot analysis

[*PSI*$^+$] *colony color assay.* The 74D-694 contains a *ade1-14* [*PSI*$^+$] suppressible nonsense allele[5]. Strains plated on YPD plates after treatment or chaperone overexpression were incubated at 30°C for 2–3 days and then incubated at room temperature for 4–7 days for color development.

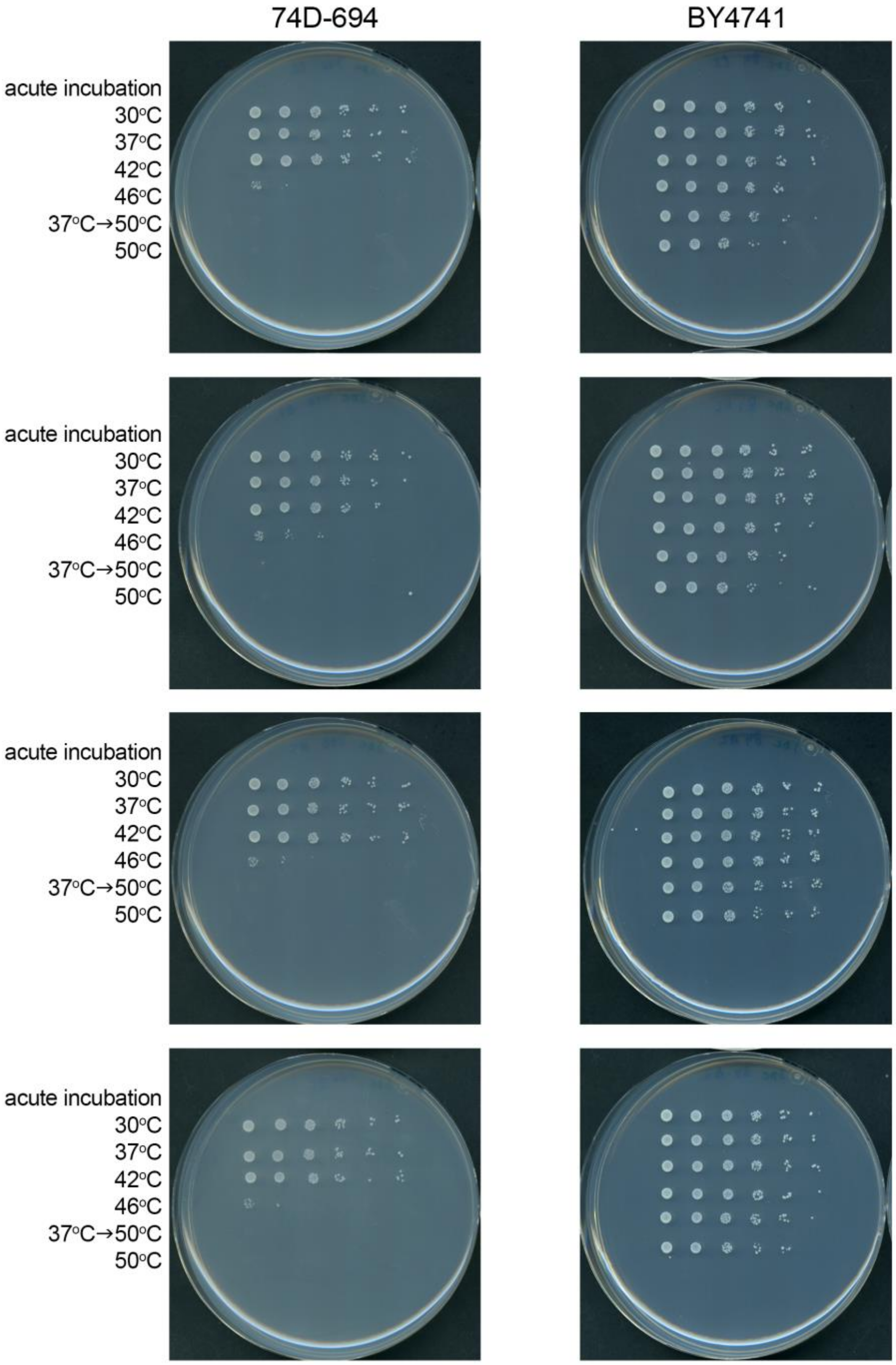


Supplemental raw data: Acute heat shock of 74D-694 and BY4741 strains. Relevant to Figure S1A.

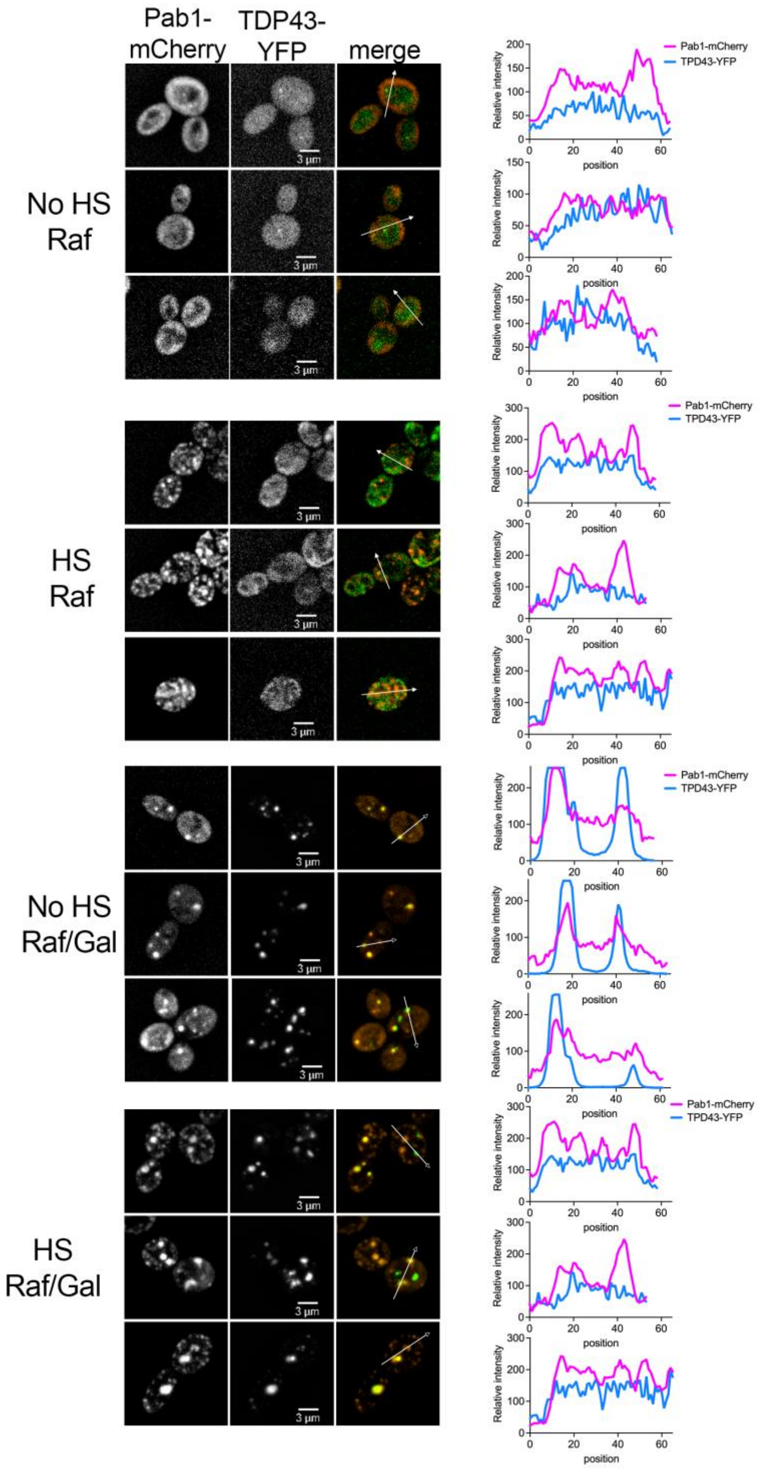


Supplemental raw data: Multiple trials co-localization images of Pab1-mCherry and TDP-43-YFP under Raf or Raf/Gal and with and without heat shock. Relevant to Figure 1A.

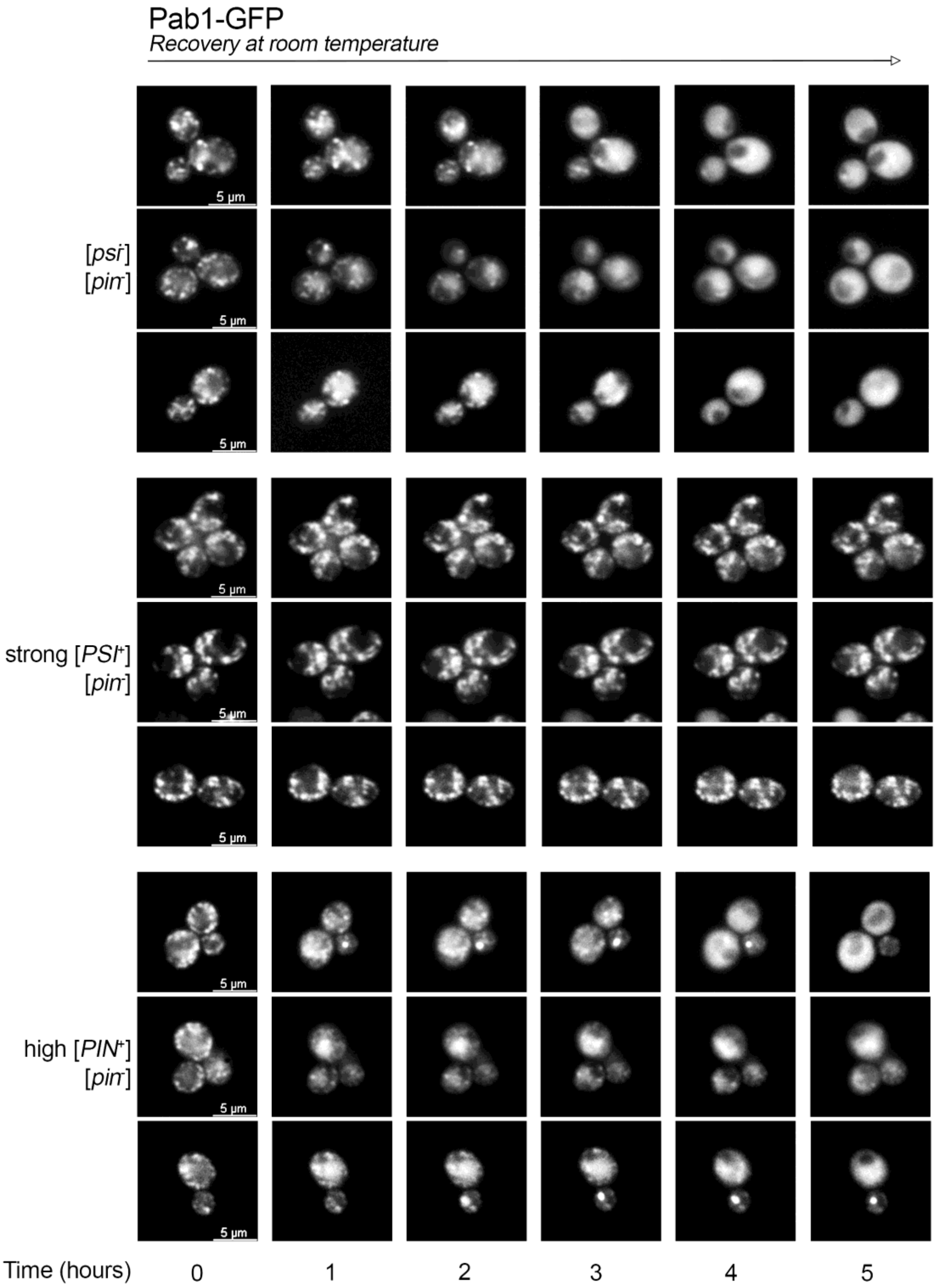


Supplemental raw data. Multiple trials timelapse of Pab1-GFP with and without prions after heat shock treatment. Relevant to Figure 2A.

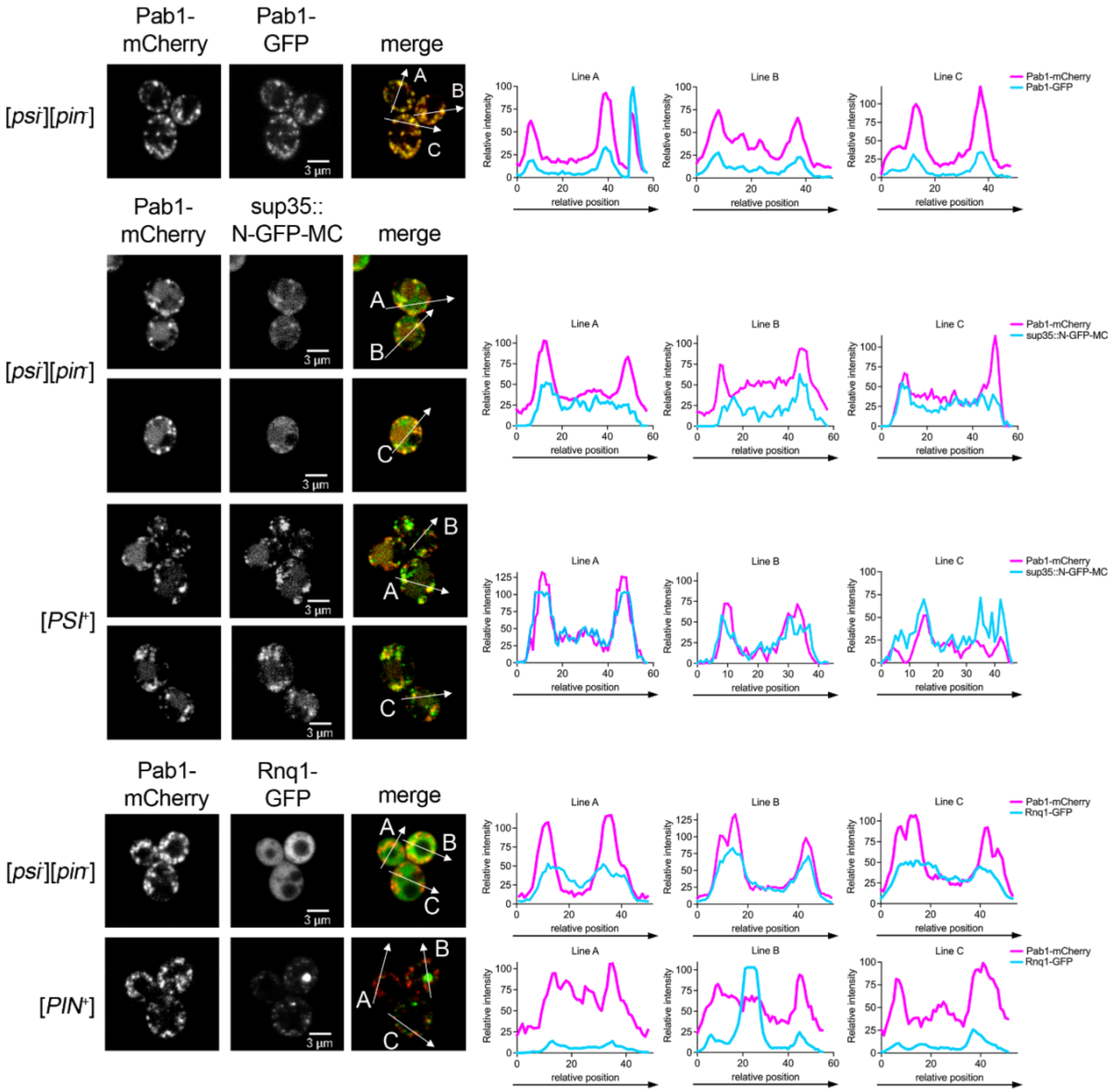


Supplemental raw data. Multiple trials of co-localization of no prion, and prion strains immediately after heat shock. Relevant to Figure 3A and B.

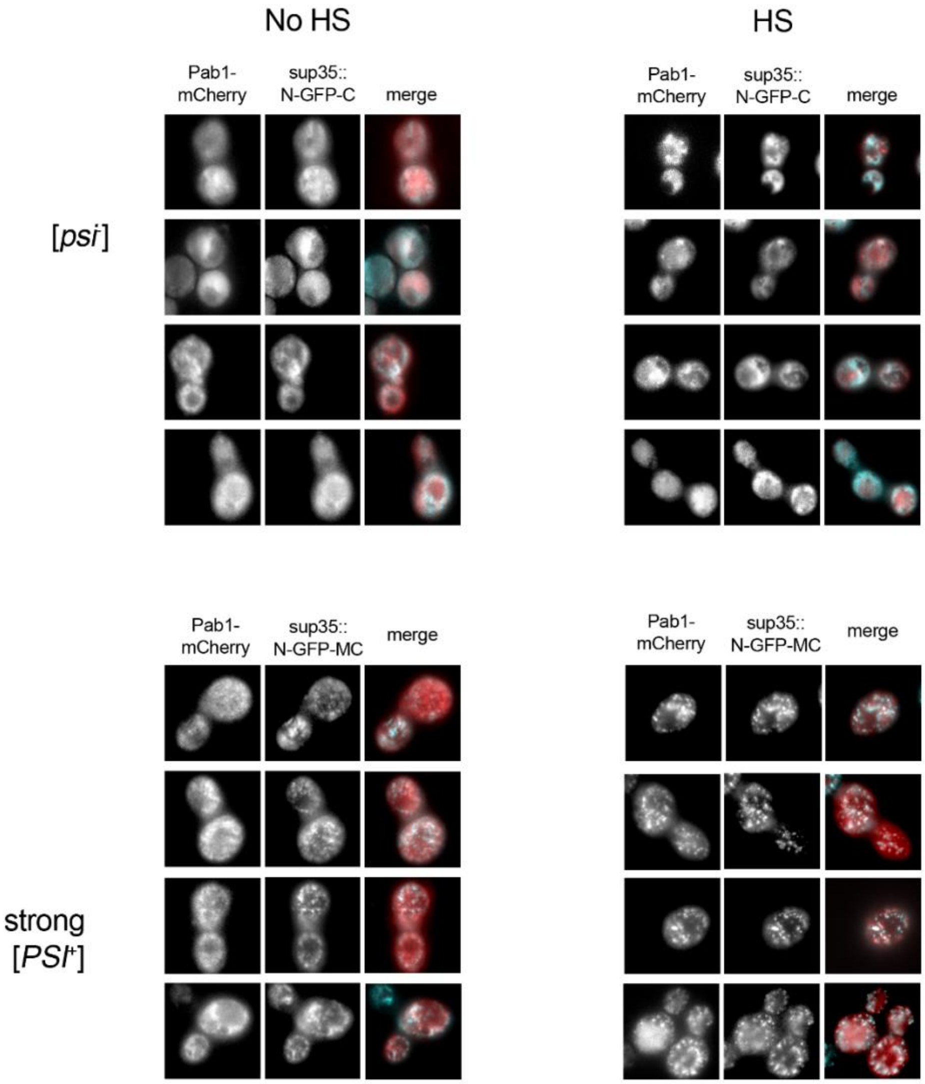


Supplemental raw data. Multiple trials of near TIRF co-localization of no prion and [*PSI*+] strains before and immediately after heat shock. Relevant to Figure 3A.

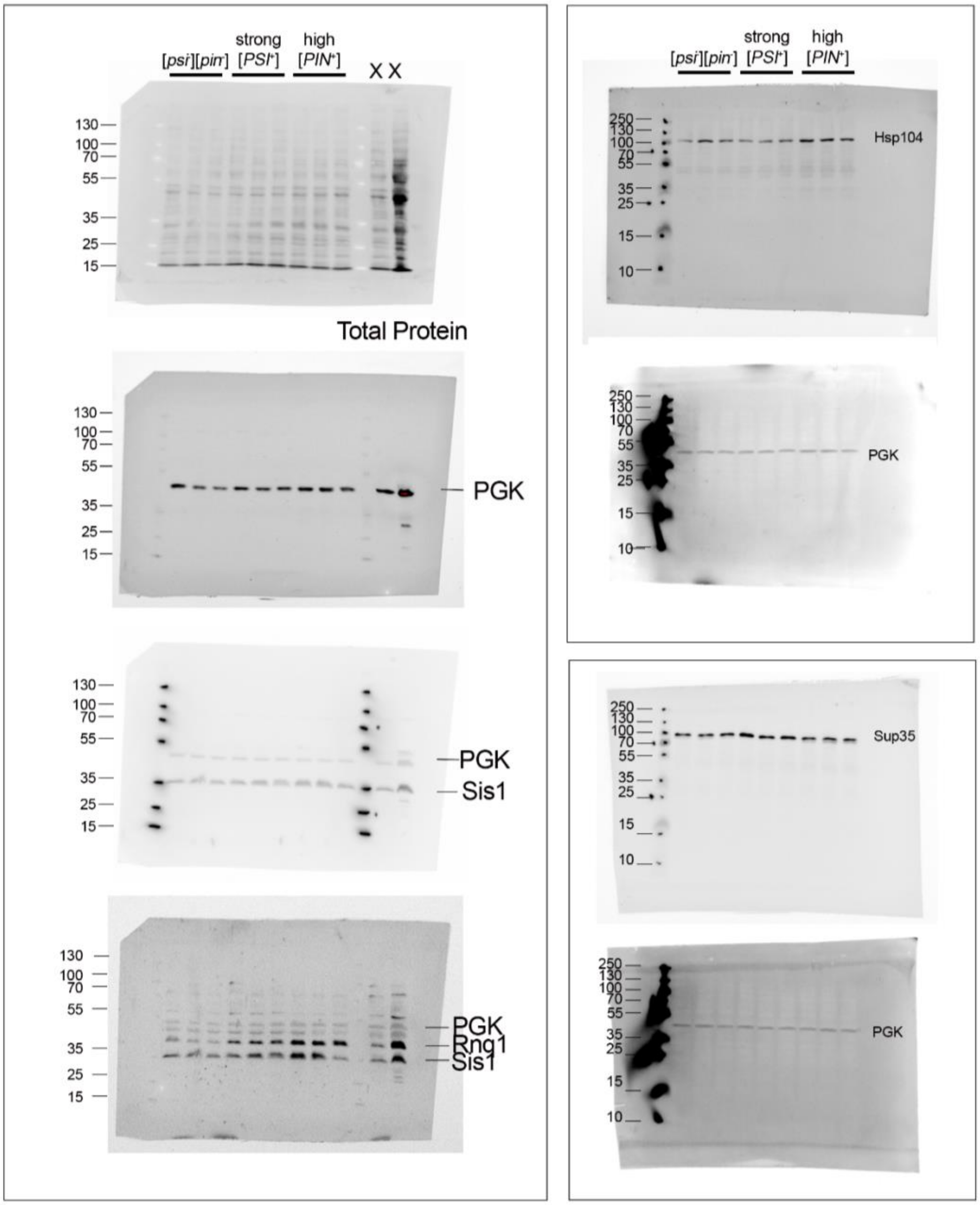


Supplemental raw data. Raw blots of steady state levels of the indicated proteins. Relevant to Figure S5B.

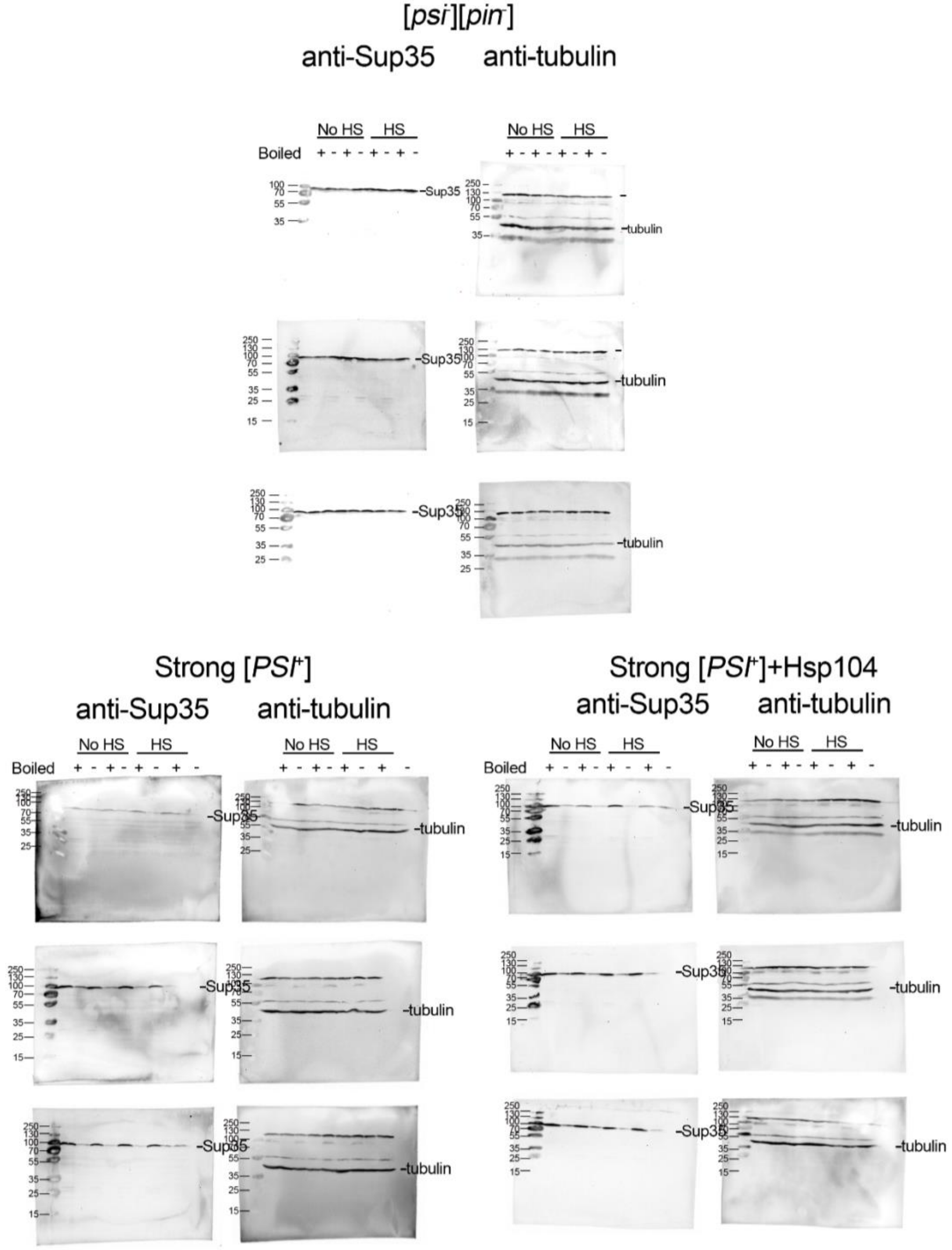


Supplemental raw data. Raw blots of well trap assays for no prion and [*PSI*+] strains. Relevant to Figure S7A.

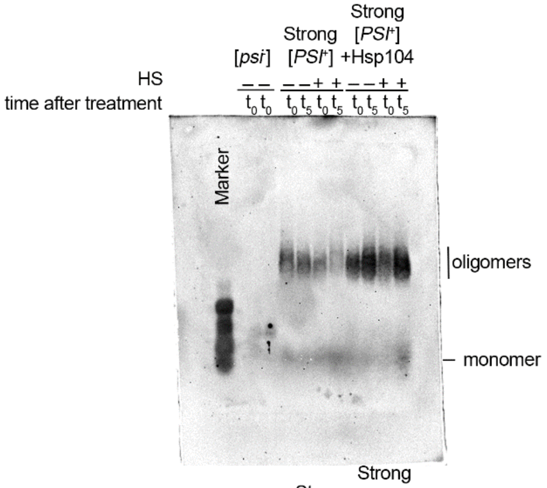


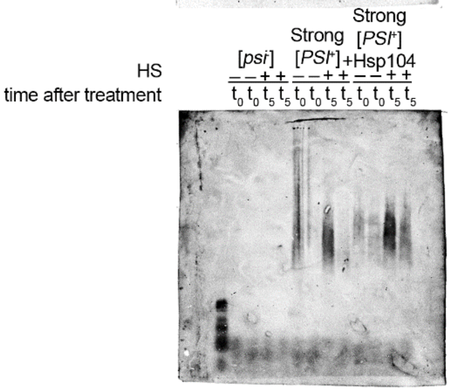


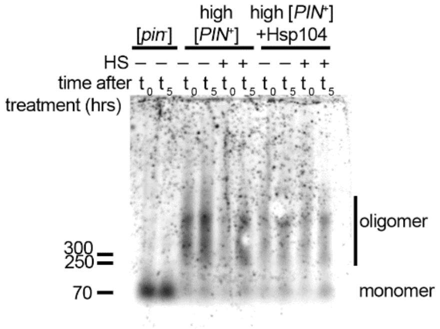


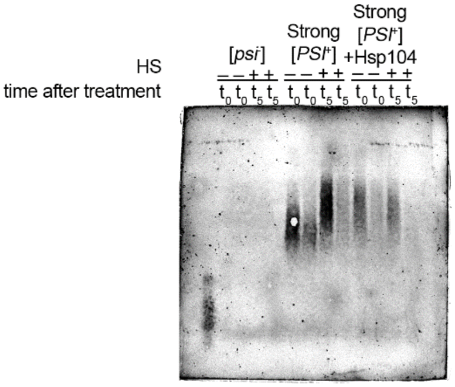


Supplemental raw data. Raw blots of SDD-AGE for no prion and prion strains. Relevant to Figure S7A and B.

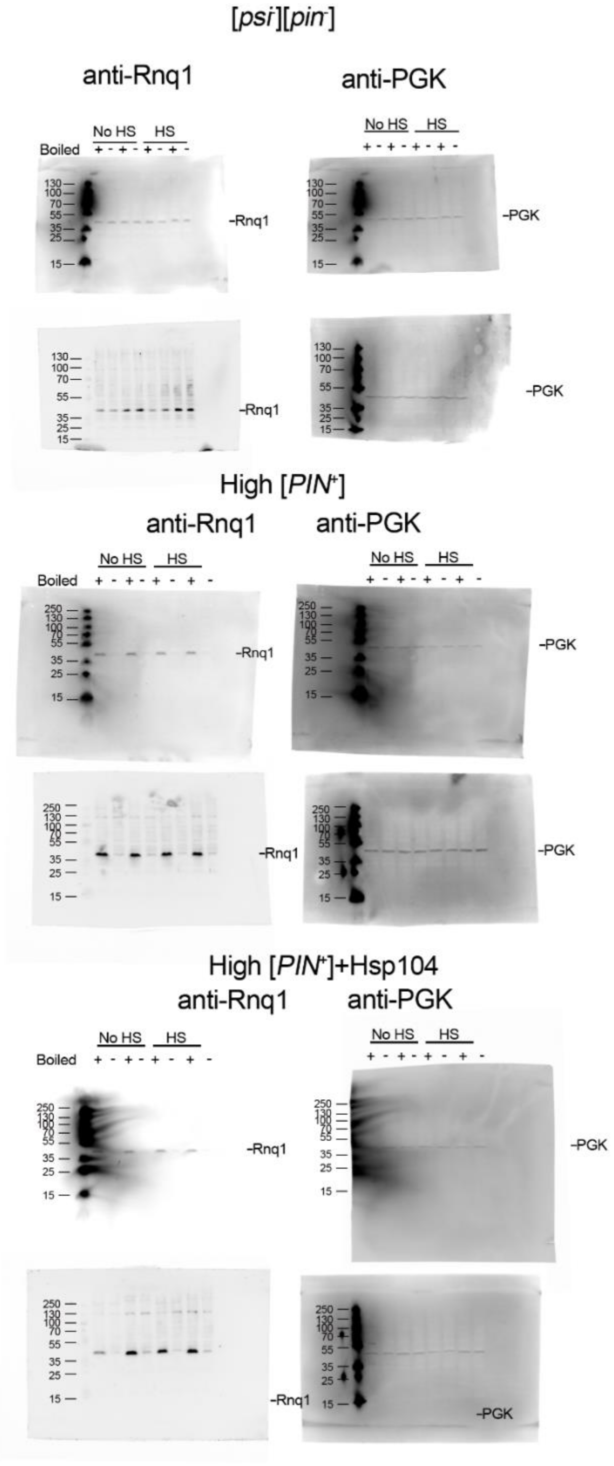


Supplemental raw data. Raw blots of well trap assays for no prion and [*PIN*$^+$] strains. Relevant to Figure S7B.

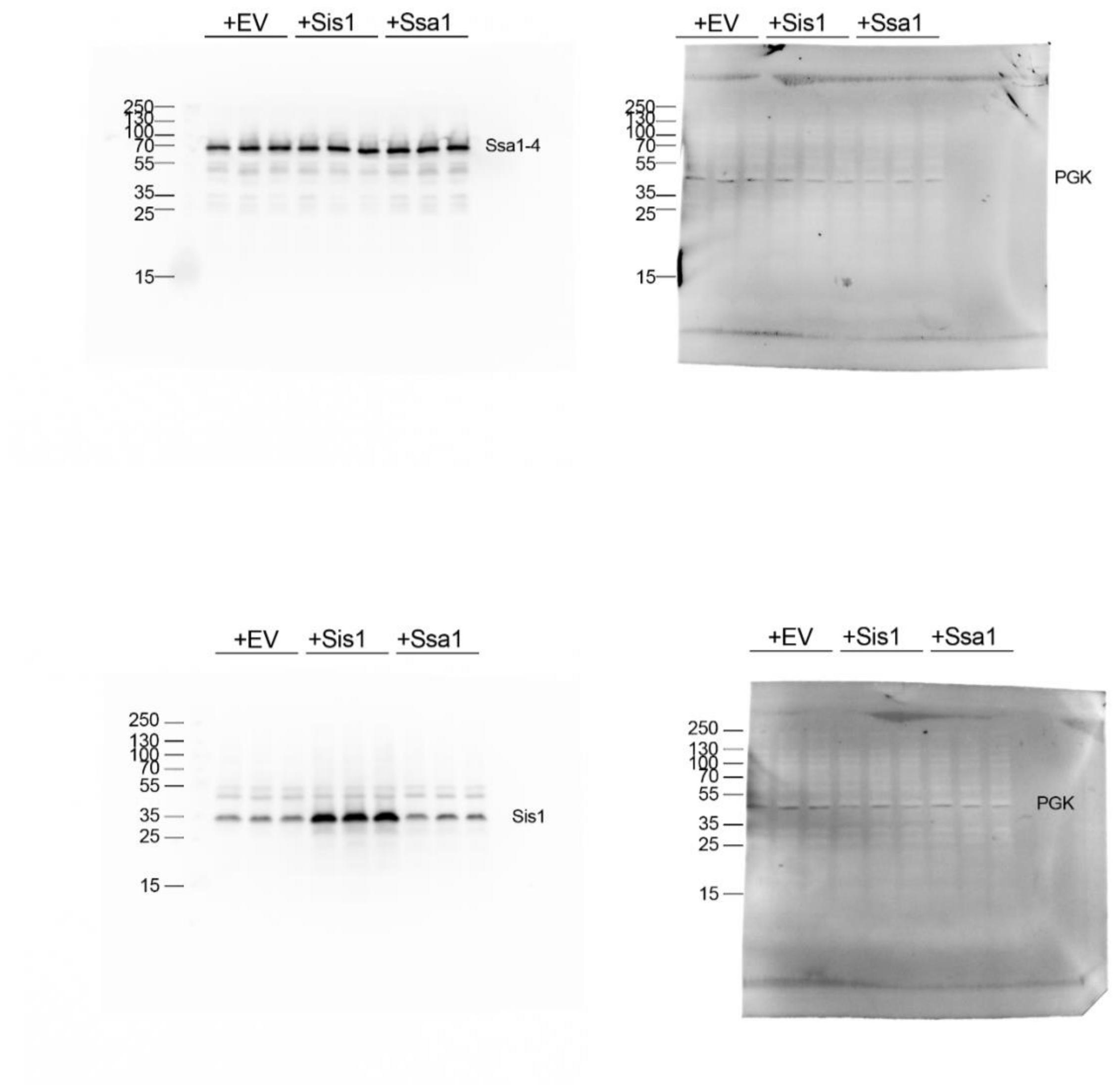


Supplemental raw data. Raw blots of steady state levels of the indicated proteins with Sis1 and Ssa1 overexpression. Relevant to Figure S8B.

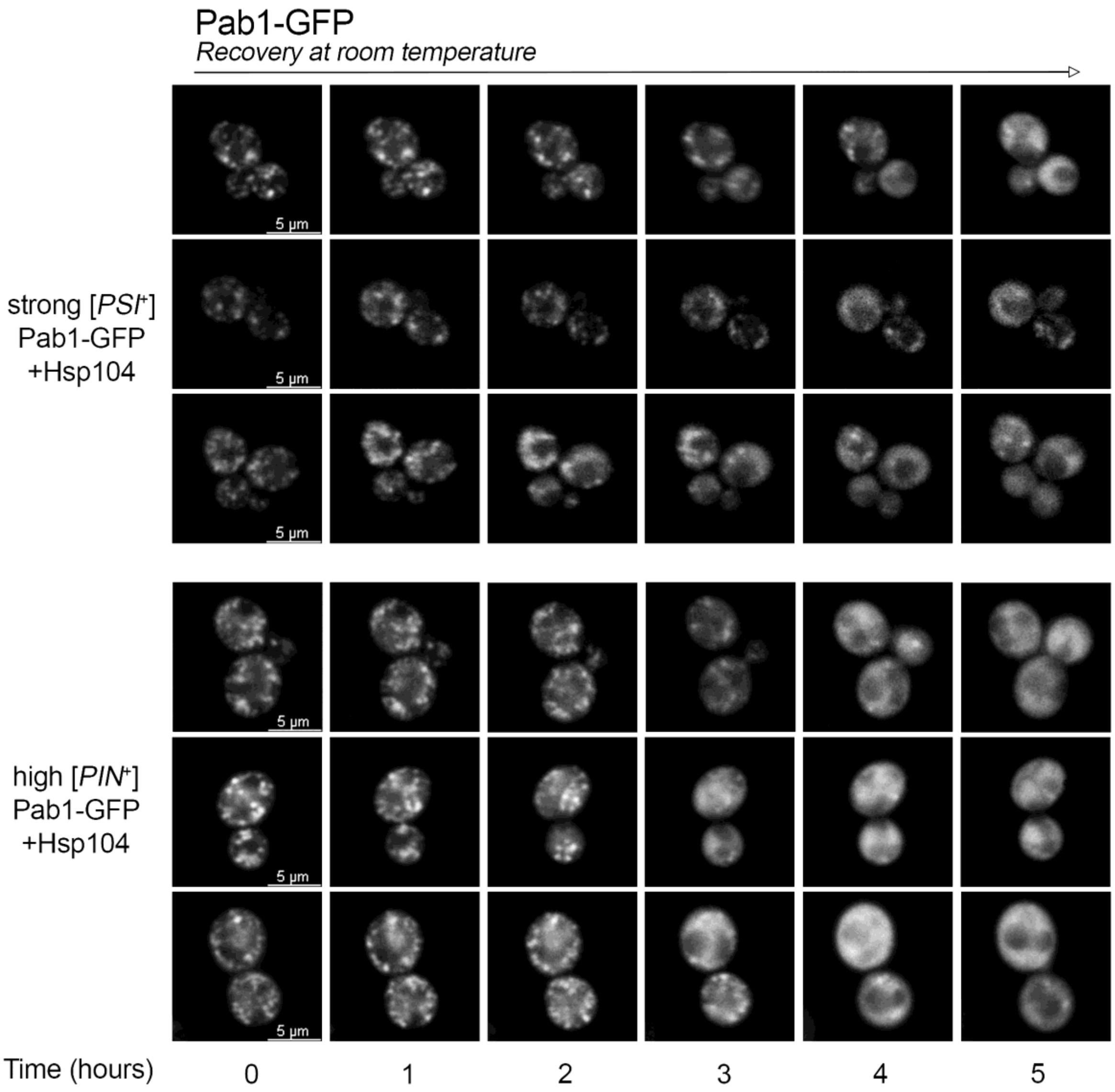


Supplemental raw data. Multiple trials of timelapse with Hsp104 overexpression of Pab1-GFP with and without prions after heat shock treatment. Relevant to Figure 4A.
